# Extracellular Matrix Mechanobiology in Pancreatic Ductal Adenocarcinoma: Correlating In Vivo Patient Magnetic Resonance Elastography with Ex Vivo Tissue Mechanics and Histopathology

**DOI:** 10.64898/2026.06.07.730664

**Authors:** Oihane Mitxelena-Iribarren, Daniela S. Garske, Dag Wulsten, Iñigo Mendizabal-Arrieta, Kristin Spirgath, Salma Almutawakel, Rosa B. Schmuck, Ingolf Sack, Amaia Cipitria

**Affiliations:** Group of Bioengineering in Regeneration and Cancer, Biogipuzkoa Health Research Institute, Donostia-San Sebastián 20014, Spain; Julius Wolff Institute, Berlin Institute of Health at Charité – Universitätsmedizin Berlin, Berlin 13353, Germany; Vicomtech Foundation, Basque Research and Technology Alliance (BRTA), Donostia-San Sebastián 20009, Spain; Institute of Medical Informatics, Charité – Universitätsmedizin Berlin, Corporate Member of Freie Universität Berlin and Humboldt-Universität zu Berlin, Berlin, Germany; Department of Radiology, Charité – Universitätsmedizin Berlin, Corporate Member of Freie Universität Berlin and Humboldt-Universität zu Berlin, Berlin, Germany; Department of Surgery, Charité – Universitätsmedizin Berlin, Campus Charité Mitte, Campus Virchow-Klinikum, Berlin 10117, Germany; IKERBASQUE, Basque Foundation for Science, Bilbao 48009, Spain

**Keywords:** Human pancreatic ductal adenocarcinoma (PDAC), in vivo magnetic resonance elastography (MRE), fresh tissue biopsy, ex vivo biomechanical testing, stiffness, viscosity, histopathology, image-based multimodal analysis

## Abstract

Pancreatic ductal adenocarcinoma (PDAC) is characterized by a dense desmoplastic extracellular matrix (ECM) that contributes to tumor progression, therapeutic resistance, and poor patient survival. However, the relationship between in vivo imaging-derived mechanical properties, ex vivo tissue biomechanics, ECM architecture, and cellularity remains incompletely understood. Here, we combined pre-operative in vivo clinical magnetic resonance elastography (MRE) with ex vivo biomechanical testing of fresh human PDAC tissue and histopathological analyses. Nine patients undergoing pancreatic resection were prospectively enrolled. Quantitative MRE was performed pre-operatively to assess tissue stiffness through shear wave speed (c) and relative viscosity or fluidity through the loss angle (φ). Fresh tumor and adjacent non-malignant tissue biopsies were subsequently analyzed ex vivo by unconfined uniaxial compression testing to determine elastic moduli and stress relaxation halftime. Histological analyses quantified collagen-rich fibrous tissue area, cell nuclei density, and nuclear morphology. Tumor tissue exhibited significantly increased stiffness and collagen fraction compared with adjacent non-malignant tissue, together with reduced cellularity, smaller nuclear area and more elongated nuclei. Ex vivo stiffness positively correlated with collagen content and negatively correlated with patient survival. Reduced stress relaxation halftime, indicative of increased tissue viscosity, was associated with lower cellularity and elongated nuclei. Importantly, pre-operative MRE parameters of the intact surrounding environment correlated significantly with ex vivo tumor mechanics, cellular organization, and survival. Specifically, a softer and less viscous surrounding environment was associated with stiffer and more viscous tumors, with lower cellularity and elongated nuclei, and poorer prognosis. These findings demonstrate that MRE-derived mechanical biomarkers reflect underlying ECM remodeling and tumor mechanobiology in PDAC. Integrating in vivo imaging with ex vivo tissue mechanics and histopathology may improve non-invasive disease characterization and support biomechanically-informed therapeutic strategies.

## 1. Introduction

Pancreatic Ductal Adenocarcinoma (PDAC) is a highly aggressive tumor characterized by therapeutic resistance and poor prognosis. A dense desmoplastic stromal microenvironment plays a central role in disease progression and treatment failure. Accounting for approximately 85–90% of pancreatic cancers, PDAC remains one of the leading causes of cancer-related mortality worldwide, with limited improvement in long-term survival, despite advances in systemic therapy and surgical management [1]. The 5-year survival rate for PDAC remains at 13% (American Cancer Society 2016) and it varies between <1% to up to 17% depending on whether patients can undergo surgical resection or not [2]. Ongoing research is focused on improving early detection, understanding tumor biology, and developing more effective personalized therapeutic strategies.

A defining pathological feature of PDAC is the extensive remodeling of the extracellular matrix (ECM), which contributes to a rich fibrotic stroma production, increased tissue stiffness, impaired vascular perfusion, immune evasion and impaired drug delivery and efficacy [3]. Increasing evidence suggests that the biomechanical properties of the tumor microenvironment are closely linked to tumor aggressiveness, cellular organization, and therapeutic response [4–8]. Various studies report biomechanical properties of human PDAC. Kpeglo and colleagues refer to values of >1 kPa for fibrotic stroma [4]. Rice et al use atomic force microscopy to measure changes in mechanical properties of healthy to PDAC in a mouse model, reporting values of 1 kPa for healthy and 4 kPa for PDAC [5]. Rubiano et al use an indentation-based method to compare the steady state modulus (kPa), viscosity (kPa s) and stress relaxation halftime (s) of healthy and PDAC in human samples [6]. They report a steady state modulus of 1.06 ± 0.25 kPa for healthy vs. 5.46 ± 3.18 kPa for PDAC; a viscosity of 252 ± 134 kPa s for healthy vs. 349 ± 222 kPa s for PDAC; and a stress relaxation halftime of 92.7 ± 46.4 s for healthy vs. 66.1 ± 20.8 s for PDAC [6]. Peerani et al use an indentation-based method to compare the elastic modulus of PDAC vs. adjacent non-malignant tissue in human samples, reporting values of 2.2 ± 0.2 kPa for non-malignant vs. 7.4 ± 0.6 kPa for PDAC [8]. However, mechanical properties of fresh human PDAC tissue, including stiffness and viscosity of tumor tissue and adjacent non-malignant tissue, have not been characterized in a systematic manner, and reported values vary depending on sample size, preparation and testing methods used [7]. Furthermore, the interplay between ECM mechanics, tissue architecture and cellularity remain incompletely understood.

Magnetic resonance elastography (MRE) has emerged as a promising non-invasive imaging modality capable of quantifying tissue mechanical properties in vivo. Tissue stiffness and viscosity can be determined with high accuracy using MRE [9]. In the context of pancreatic human tissue, Zhu et al [10] measured the shear wave speed (c, in m/s) as a surrogate marker of stiffness and the loss angle ((*φ*, in rad) as a proxy for relative viscosity or tissue fluidity [11]. They reported values of 1.32 ± 0.05 m/s for surrounding non-tumor environment vs. 2.35 ± 0.39 m/s for PDAC tumor, and 0.81 ± 0.04 rad for non-tumor environment vs. 1.20 ± 0.13 rad for PDAC tumor [10], with higher stiffness and viscosity for tumor tissue. Marticorena et al also employed MRE to measure c in human pancreas, reporting values of 1.28 ± 0.14 m/s for non-tumor environment vs. 2.08 ± 0.38 m/s for PDAC tumor [12]. Schattenfroh et al employed MRE with healthy volunteers and reported values for human pancreas of c 1.31± 0.12 m/s [13]. However, the relationship between patient-derived MRE measurements and the underlying ex vivo tissue biomechanics, ECM composition, architecture, and cellularity in PDAC remains poorly understood. Correlative studies integrating radiologic, mechanical, and histopathological analyses are therefore essential to elucidate how macroscopic imaging biomarkers reflect the complex biophysical characteristics of the tumor tissue and surrounding environment.

The aim of this study is to correlate pre-operative in vivo clinical MRE imaging of patients, including measurements of stiffness and tissue viscosity, with ex vivo fresh tissue biopsy biomechanical testing, ECM tissue architecture, cell nuclei density and morphology analysis, in order to improve our understanding how ECM mechanical properties modulate disease progression. Improved understanding of these relationships may enhance non-invasive disease characterization with biophysical properties, facilitate patient stratification, and support the development of biomechanically informed therapeutic approaches in PDAC.

## 2. Materials and methods

### 2.1 Patient sample collection and clinical evaluation

All patients underwent partial pancreatic resection at the Department of Surgery, Charité – Universitätsmedizin Berlin between 2018 and 2020. Written informed consent was obtained from all participants prior to inclusion. The study protocol was approved by the institutional ethics committee (Ethikkommission der Charité, EA1/188/17, EA1/076/17) and conducted in accordance with the principles of the Declaration of Helsinki and its later amendments.

Prior to surgery, all patients underwent magnetic resonance imaging (MRI) examination for posterior quantitative magnetic resonance elastography (MRE) (see section 2.2). Depending on the underlying pathology and tumor location, patients underwent either pancreatic head resection in the form of pylorus-preserving pancreaticoduodenectomy (PPPD) or distal pancreatectomy (DP), including left-sided resections and Appleby procedures.

Following pancreatic head resection, reconstruction of the pancreatic remnant was carried out either by pancreatogastrostomy or pancreatojejunostomy. In distal pancreatectomy, stump closure was performed either using a Seamguard-reinforced stapled transection with a 60 mm Covidien stapler or by a bovine serum albumin–glutaraldehyde sealed fish-mouth technique as previously described [14]. In all cases, an intraoperatively placed drain was positioned adjacent to the pancreatic anastomosis or remnant and maintained for a minimum of four postoperative days. Surgical management and perioperative care followed previously established institutional protocols [15]. All procedures were performed by experienced pancreatic surgeons with an annual case volume exceeding 50 pancreatic resections.

Patients were monitored clinically until hospital discharge. Recorded perioperative parameters included neoadjuvant therapy, relevant medical history such as pancreatitis or cholestasis, postoperative outcomes including drain fluid analysis, and routine laboratory findings. POPF was diagnosed according to the criteria of the International Study Group of Pancreatic Surgery (ISGPS), defined as drain fluid lipase levels exceeding three times the upper normal serum value on or after postoperative day three [16].

Histopathological evaluation of pancreatitis was performed on tissue adjacent to the resection margin. Cholestasis was defined by elevated alkaline phosphatase (AP) and gamma-glutamyltransferase (GGT) levels greater than 2.5-fold above the upper reference limit and/or radiologic evidence of significant biliary obstruction. Dilatation thresholds were defined as follows: pancreatic duct diameter > 3 mm and bile duct diameter > 6 mm, or > 10 mm in patients after cholecystectomy.

Immediately after resection, the surgical specimen was assessed by a pathologist. Tissue samples were also obtained from non-tumorous pancreatic tissue adjacent to the resection margin. For biomechanical analysis, tissue cylinders were collected using a standardized 5 mm Kai biopsy punch (PFM Medical, Cologne, Germany), modified with a custom plunger system to improve specimen retrieval [17]. Mechanical testing of the fresh tissue tumor and non-malignant samples was carried out ex vivo within two hours following resection, as in previous studies [17].

### 2.2 Pre-operative in vivo clinical magnetic resonance elastography (MRE) imaging of patients

All quantitative magnetic resonance imaging (MRI) examinations were performed on a 3-T MRI system (Magnetom Vida, Siemens, Erlangen, Germany). Images were obtained in transversal slice orientation. Diffusion weighted imaging (DWI) was performed with 104 *×* 128 *×* 11 matrix size and 2.7 *×* 2.7 *×* 5 mm³ voxel size. b-values of 0 and 500 s/mm² were acquired to compute apparent diffusion coefficient (ADC) maps (in µm²/s). MRI volumetry was based on 3D T2-weigthed MRI with 192 *×* 256 *×* 23 matrix size and 1.6 *×* 1.6 *×* 7 mm³ voxel size. PDAC volumes were measured by manual segmentation in 3D. Multifrequency MRE was performed using a spin-echo echo-planar imaging sequence [18,19] acquired at external vibration frequencies of 30, 40, 50 and 60 Hz induced by four compressed air drivers fixed to the chest with Velcro strips (Vibro4.2, THEA devices, Wurzen, Germany). MRE matrix size was 104 x 128 x 25 with 2.7 x 2.7 x 5 mm³ voxel size.

MRE postprocessing was performed using a server-based software (https://bioqic-apps.charite.de) [20], which provides implementations of wavenumber-based (k-) multifrequency elasto-visco inversion (k-MDEV) for reconstruction of shear wave speed (c) [21] and MDEV for reconstruction of the loss angle ((*φ*) of the complex shear modulus [22]. c (in m/s) was determined as a surrogate marker of stiffness, while (*φ* (in rad) was used as a proxy for relative viscosity or tissue fluidity [23]. Fluidity of solid tissue is related to the slope of c dispersion over frequency [9]. c and *ϕ* are calculated both as average of the PDAC tumor (tumor) and as average of the surrounding environment (env).

### 2.3 Ex vivo mechanical testing of human tissue biopsies

The mechanical properties of fresh PDAC tissue biopsies were determined by ex vivo unconfined uniaxial compression testing (Test Bench LM 1 ElectroForce, Bose, Eden Prairie, MN). Mechanical testing was conducted within 2 h after tissue resection, in air at room temperature, as described previously [17]. Briefly, the samples were compressed at a rate of 1 mm/min and were not additionally hydrated for the duration of the experiment to avoid any material alterations. Load (50 lbs / 225 N load cell) and displacement data were sampled at 100 Hz. In order to convert these into stress and strain values, the area of the tissue biopsy was defined by the 5 mm diameter of the biopsy punch and the initial height of the sample was determined by the distance between the parallel plates upon first contact at a preload of 0.2 g.

The tissue stiffness was measured by the elastic modulus, E (kPa), which was calculated based on the best-fit slope of the stress σ (kPa) vs. strain ε (%) curves, in both the linear (E_lin, Pa) and non-linear elastic range (E_non lin, Pa), using a routine written in MatLab R2019b (The Mathworks, Inc. Natick, MA, USA) (see Data Availability). To measure the stress relaxation at 15 % strain the displacement was kept constant for approximately 500 seconds. The stress relaxation halftime, τ_1/2_, (s), was calculated as the time interval between the maximum stress and the time needed to drop to half its value, at constant 15% strain. After mechanical testing, samples were fixed in 4% paraformaldehyde for 2 h, at 4°C. Samples were stored in PBS at 4°C until cryoembedding.

### 2.3 Histology and polarized light microscopy

Prior to histological analyses, samples underwent gradual dehydration in ascending sucrose solutions (10 %, 20 % and 30 % sugar), incubating for 24 h at 4°C each step. They were subsequently embedded in Tissue-Tek O.C.T. compound (SAKURA, #4583). 7 μm thick cryosections were cut with a cryotome (Cryostar NX70, Thermoscientific), dried for 60 min at RT and stored at -80°C. Prior to staining, the frozen sections were dried for 45 minutes at RT.

To analyze cell density and cell morphological characteristics, cryosections were stained with Hematoxylin (Harris-Hematoxylin) and Eosin with a differentiation step in HCl alcohol in between, and finally mounted in Vitroclud (R. Langenbrinck GmbH, #04-0001). Images were acquired with an optical microscope (DM6B, Leica) in brightfield mode with 40x magnification. For image analysis, a custom-made Python code (version 3.10.11) was used (see Data Availability). First, the total tissue area was defined by thresholding the void areas and other cutting artefacts. Then the images were cut into 5000 x 5000 pixel (2095 x 2095 µm) by using the OpenSlide library (version 3.4.1) in Python. Next, using the software StarDist (version 0.8.5, https://github.com/stardist/stardist) [24–26], individual nuclei were segmented. Finally, total cell nuclei density (#/mm^2^), single cell nuclear area (*μ*m^2^), diameter (*μ*m) and aspect ratio (-) were quantified (Supplementary Fig. S1).

To quantify the fraction of collagen fibrous tissue, cryosections were stained with Picrosirius Red, adapted from [27]. Briefly, sections were stained for 1 h in 0.1 % Picrosirius Red, washed briefly in 0.5 % acetic acid, followed by dehydration, cleared in xylene and mounted with Vitroclud. Image acquisition was performed with an optical microscope (AxioObserver 7, Zeiss) in bright field mode and with linear polarized light with six different polarization angles at 10x magnification as described before [28]. Based on the bright-field images, the total tissue area (excluding void areas) was defined. Next, using the six polarized light images of each sample, the collagen fibrous tissue area per total tissue area (%) was quantified with a newly developed Python code (Supplementary Fig. S2).

### 2.5 Statistical analysis

Statistical analysis was conducted with GraphPad Prism 10.6.1 (GraphPad Software, Inc., La Jolla, CA, USA). Distribution of variables was tested using the Shapiro-Wilkinson test. Unpaired t-test was used to test for significant differences between tumorous and non-malignant tissue. When normality distribution was not fulfilled, Mann Whitney U-test was performed. Correlations between ex vivo tissue mechanics, tissue architecture and cellularity, and pre-operative in vivo clinical MRE data were calculated applying Pearson’s or Spearman correlation. p < 0.05 determines statistically significant differences.

## 3. Results

### 3.1 Clinical evaluation

The following clinical variables are summarized on Table 1: survival of patients (evaluated from surgery date until May 2026), sex, previous pancreatitis or cholestasis, pancreatic duct dilatation, bile duct dilatation, pre-operative stenting, neoadjuvant therapy received before the surgery, type of surgery, tumor entity confirmation, Union for International Cancer Control (UICC) classification, TNM classification by Tumor size/extent, Node involvement, and distant Metastasis, postoperative pancreatic fistula (POPF) and assessment of cellularity by pathologist.

**Table 1:**
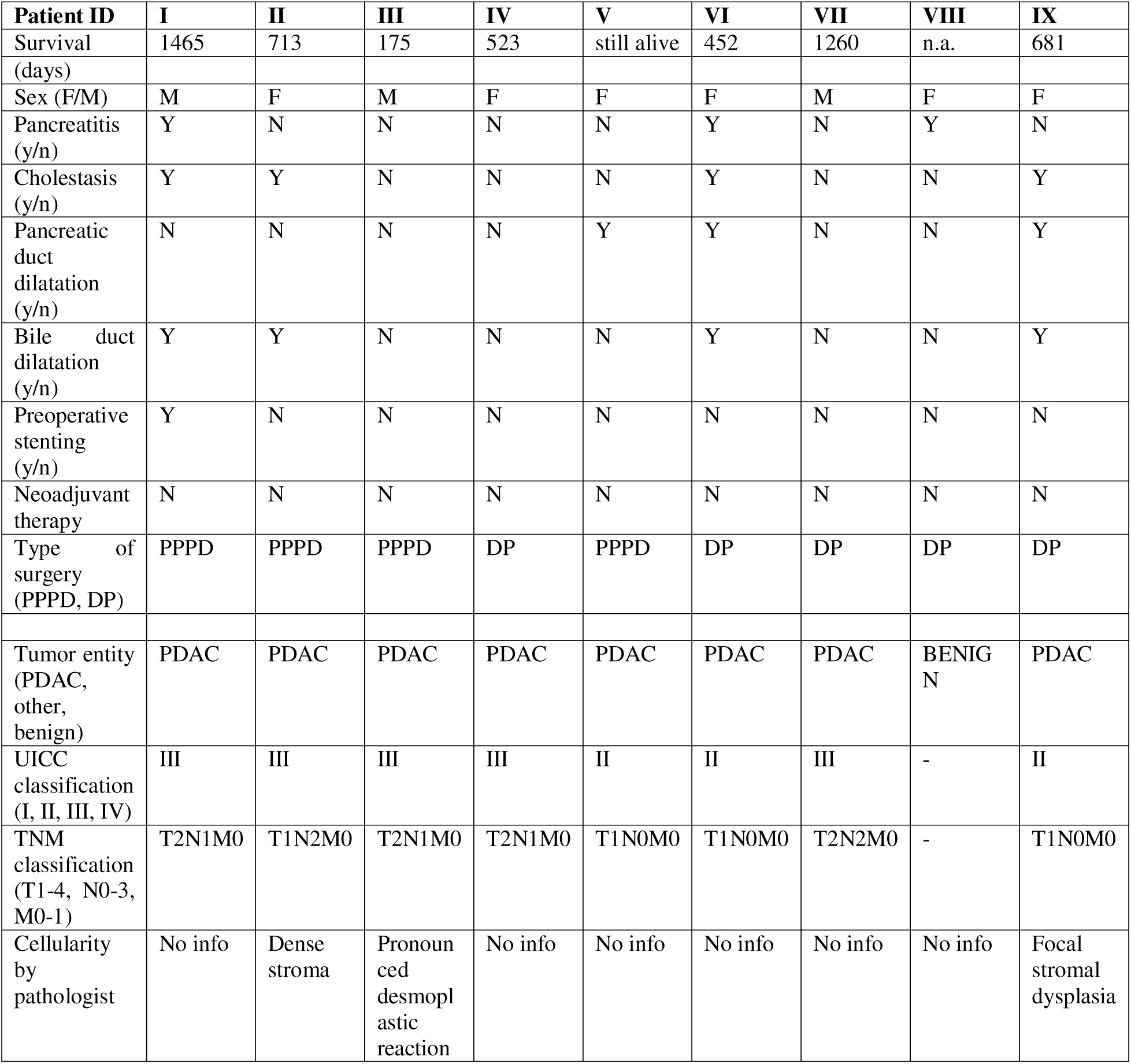
Clinical variables of the study cohort. PDAC: Pancreatic ductal adenocarcinoma, PPPD: pylorus-preserving pancreaticoduodenectomy, UICC: Union for International Cancer Control. TNM classification by Tumor size/extent, Node involvement, and distant Metastasis, POPF: postoperative pancreatic fistula. Total of 9 patients, with tumor (tumor) vs. adjacent non-malignant tissue (nm).

| Patient ID | I | II | III | IV | V | VI | VII | VIII | IX |
| --- | --- | --- | --- | --- | --- | --- | --- | --- | --- |
| Survival | 1465 | 713 | 175 | 523 | still alive | 452 | 1260 | n.a. | 681 |
| (days) |  |  |  |  |  |  |  |  |  |
| Sex (F/M) | M | F | M | F | F | F | M | F | F |
| Pancreatitis (y/n) | Y | N | N | N | N | Y | N | Y | N |
| Cholestasis (y/n) | Y | Y | N | N | N | Y | N | N | Y |
| Pancreatic duct dilatation (y/n) | N | N | N | N | Y | Y | N | N | Y |
| Bile duct dilatation (y/n) | Y | Y | N | N | N | Y | N | N | Y |
| Preoperative stenting (y/n) | Y | N | N | N | N | N | N | N | N |
| Neoadjuvant therapy | N | N | N | N | N | N | N | N | N |
| Type of surgery (PPPD, DP) | PPPD | PPPD | PPPD | DP | PPPD | DP | DP | DP | DP |
| Tumor entity (PDAC, other, benign) | PDAC | PDAC | PDAC | PDAC | PDAC | PDAC | PDAC | BENIGN | PDAC |
| UICC classification (I, II, III, IV) | III | III | III | III | II | II | III | - | II |
| TNM classification (T1-4, N0-3, M0-1) | T2N1M0 | T1N2M0 | T2N1M0 | T2N1M0 | T1N0M0 | T1N0M0 | T2N2M0 | - | T1N0M0 |
| Cellularity by pathologist | No info | Dense stroma | Pronounced desmoplastic reaction | No info | No info | No info | No info | No info | Focal stromal dysplasia |

Patients recruited for this study, after they signed the informed consent, followed the workflow described in Figure 1. Pre-operative in vivo clinical imaging was performed by MRE shortly before the surgery. After surgical resection, a pathologist evaluated the resected tissue and collected a tumor sample for this study, together with adjacent non-malignant tissue, following the procedure previously described [17]. Ex vivo mechanical testing was conducted within 2 h after tissue resection. Following this, samples were fixed and stored in PBS until further processing for histology. All histological images analyzed are collected in Supplementary Fig. S3-S11.

**Figure 1:**
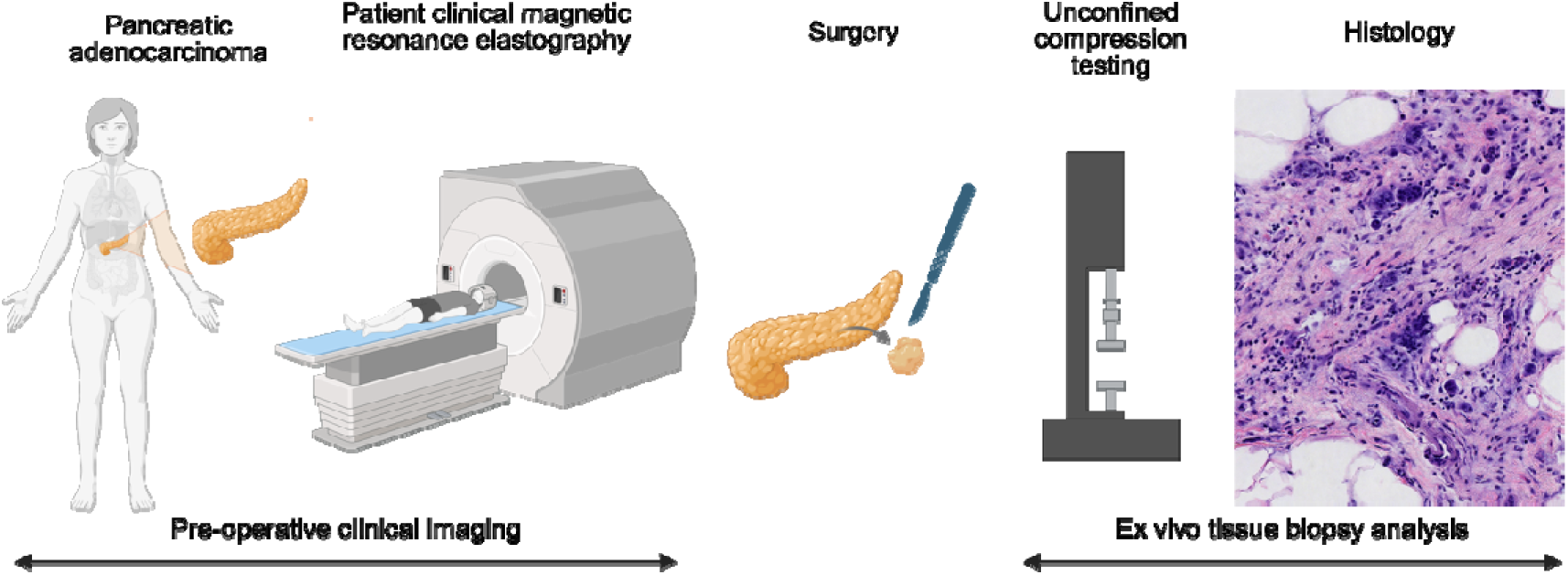
Workflow of patient pre-operative in vivo clinical imaging with MRE, followed by surgical resection, correlative ex vivo mechanical testing of fresh samples, tissue fixation, cryoembedding and histology of tissue architecture and cellularity. Created with Biorender.

### 3.2 Preoperative in vivo clinical magnetic resonance elastography (MRE) imaging of patients

Preoperative MRE was successfully performed in all patients, providing elastographic maps of the tumor and surrounding tissue before surgical intervention (Fig. 2A-D). These MRE maps revealed marked spatial heterogeneity within the tumor itself and the surrounding environment. Quantification of the shear wave speed (c) and the loss angle ( ) of the complex shear modulus was performed in both the tumor itself (tumor) and the surrounding environment (env) as a reference (Fig. 2E, F).

**Figure 2:**
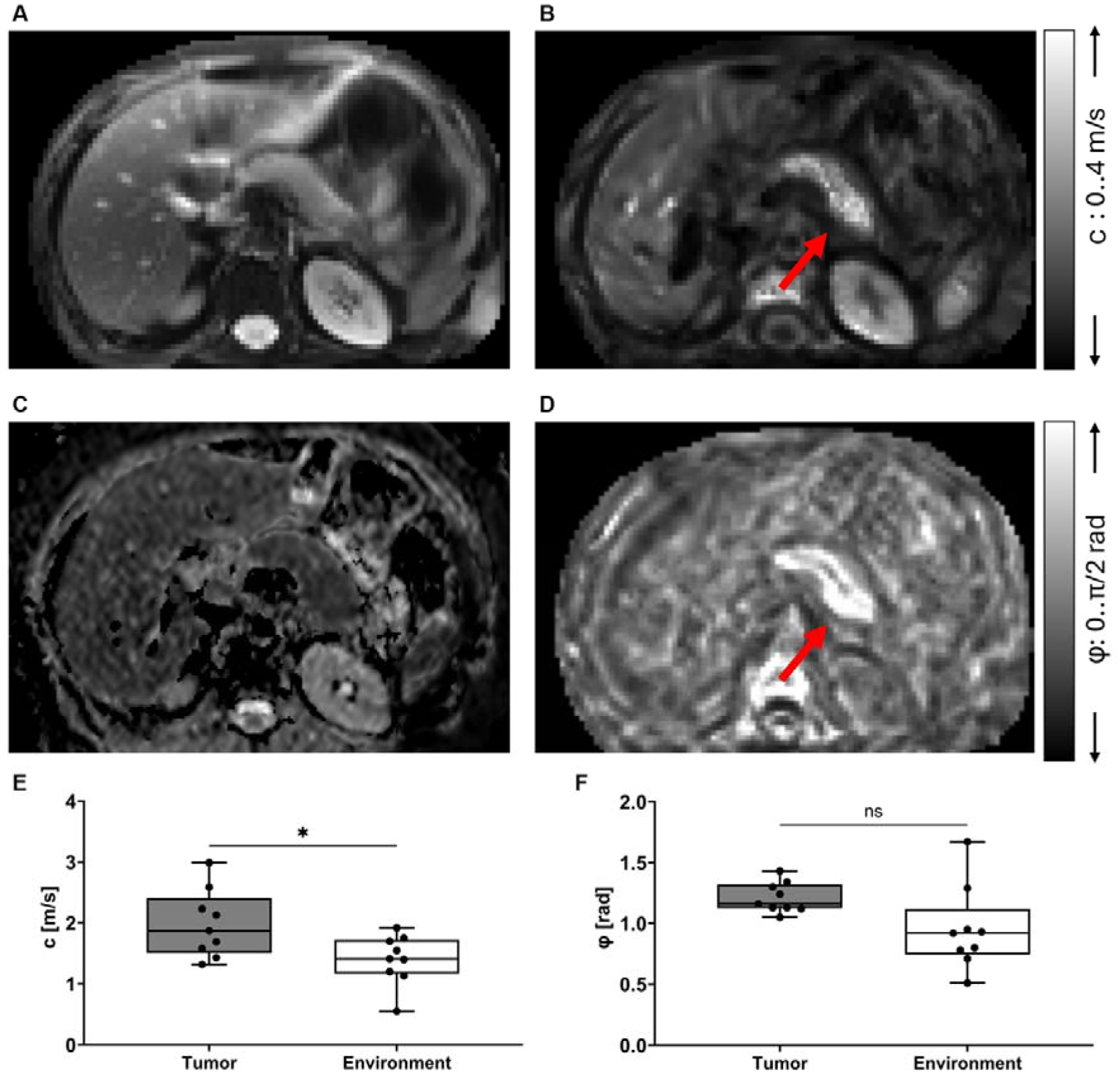
MRE maps corresponding to patient II showing the (A) MRE magnitude, (B) MRE stiffness as shear wave speed (c) map, (C) diffusion weighted imaging (DWI) of the apparent diffusion coefficient (ADC) and (D) MRE fluidity as the loss angle (φ) of the complex shear modulus. Quantification of MRE maps comparing (E) c and (F) φ values of tumor tissue vs. surrounding environment. Red arrow indicates the PDAC tumor.

Quantitative analysis showed that tumor regions exhibited a statistically increased mechanical stiffness (p = 0.024) compared with adjacent tissue, as c_tumor (1.98±0.56 m/s) was always higher than c_env (1.4±0.41 m/s). Similarly, the relative viscosity or tissue fluidity of the tumor region was found to be usually higher than its environment (although not statistically significant, p = 0.058), with a mean value of 1.21±0.12 rad and 0.95±0.34 rad respectively. ADC mean value between PDAC patients was 1239.86±116.44 µm²/s, showing consistency in this illness.

### 3.3 Ex vivo tissue mechanics, ECM architecture, cell nuclei density and morphology of human pancreatic tissue samples

The elastic and viscoelastic properties of PDAC patient tissue biopsies, comparing a tumor sample (Fig. 3) with the corresponding adjacent non-malignant tissue (Fig. 4) of the same patient, were quantified using ex vivo unconfined uniaxial compression testing of fresh samples. From the stress vs. strain curves, the elastic modulus was obtained as the slope in the linear region, E_lin, and the slope in the non-linear region, E_non lin (Fig. 3A, Fig. 4A). Both parameters were significantly higher for the tumor samples compared to the adjacent non-malignant tissue (Fig. 5A, B), with p = 0.008 and p = 0.012, respectively, confirming the stiffer nature of PDAC samples.

**Figure 3:**
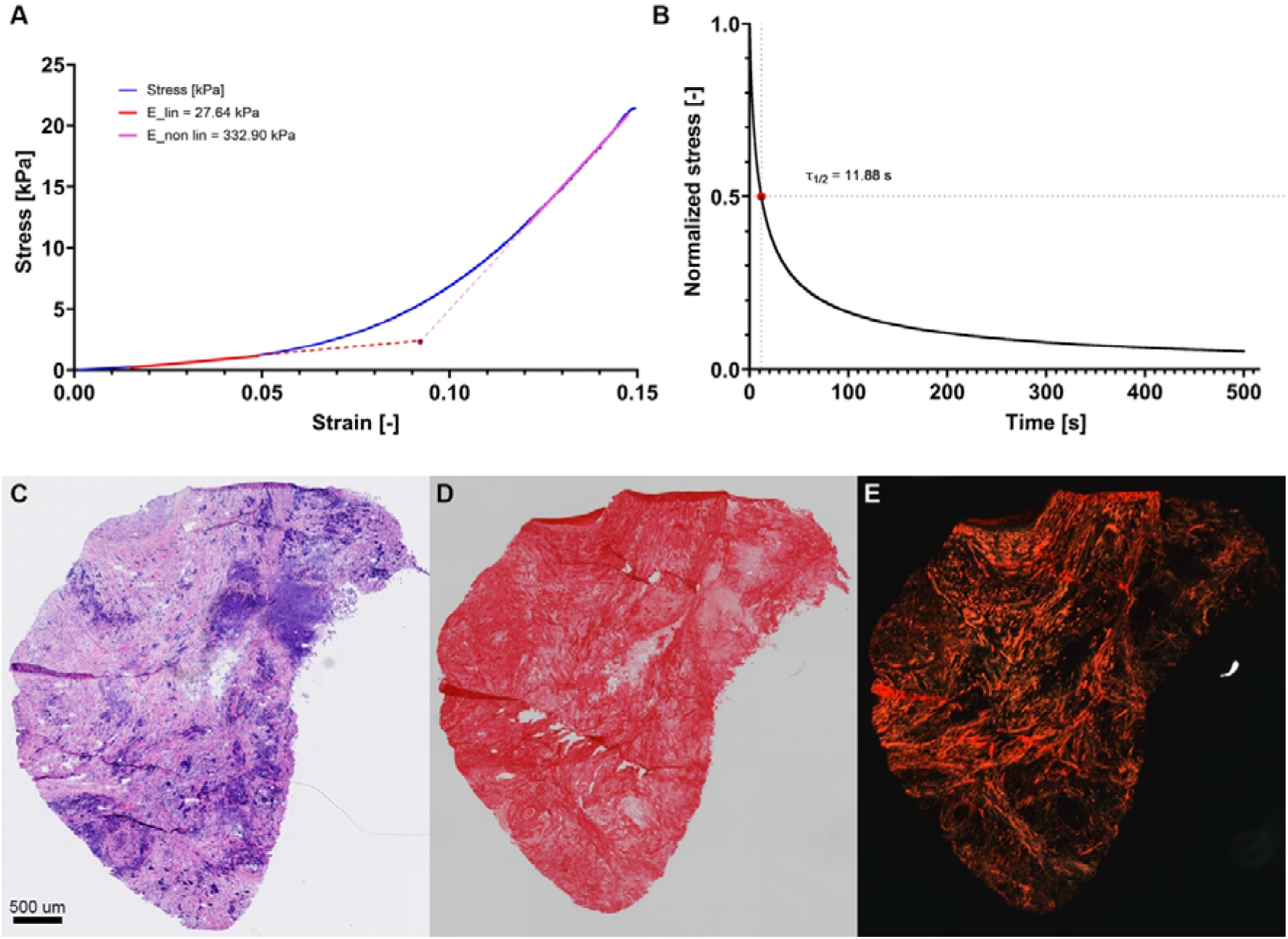
Exemplary compilation of ex vivo tissue mechanics and ECM architecture of a tumor sample (patient III) showing (A) the curve stress vs. strain in the E_lin vs. E_non lin part with corresponding slope E_lin of 26.6 kPa and E_non lin of 332.9 kPa, (B) the curve stress vs. time showing a stress relaxation halftime τ__½_ of 11.8 s, (C) H&E staining, (D) Picrosirius red staining with bright field and (E) polarized light imaging.

**Figure 4:**
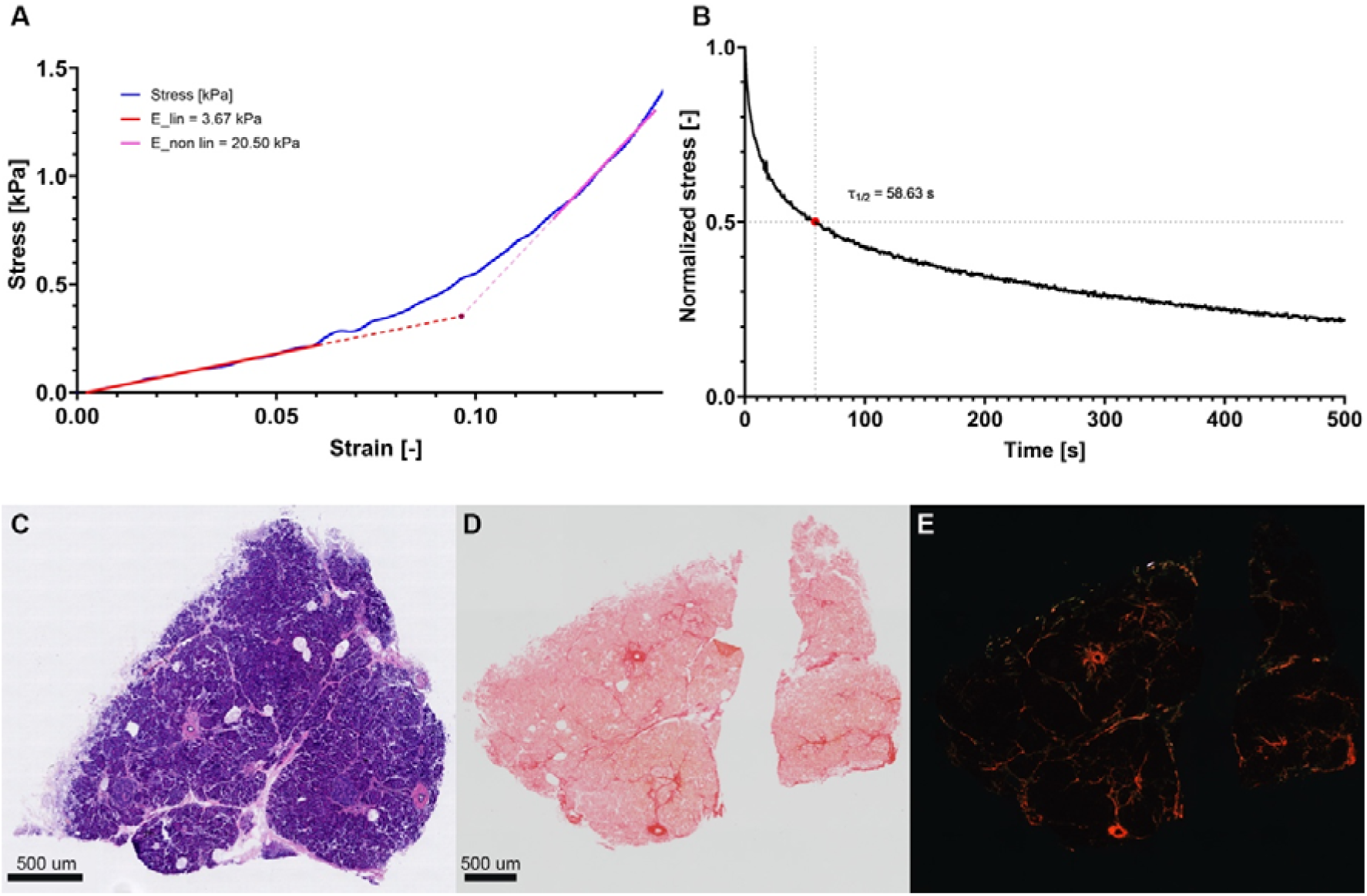
Exemplary compilation of ex vivo tissue mechanics and ECM architecture of an adjacent non-malignant tissue sample (patient III) showing (A) the curve stress vs. strain in the E_lin vs. E_non lin part with corresponding slope E_lin of 3.67 kPa and E_non lin of 20.5 kPa, (B) the curve stress vs. time showing a stress relaxation halftime τ__½_ of 58.6 s, (C) H&E staining, (D) Picrosirius red staining with bright field and (E) polarized light imaging.

**Figure 5:**
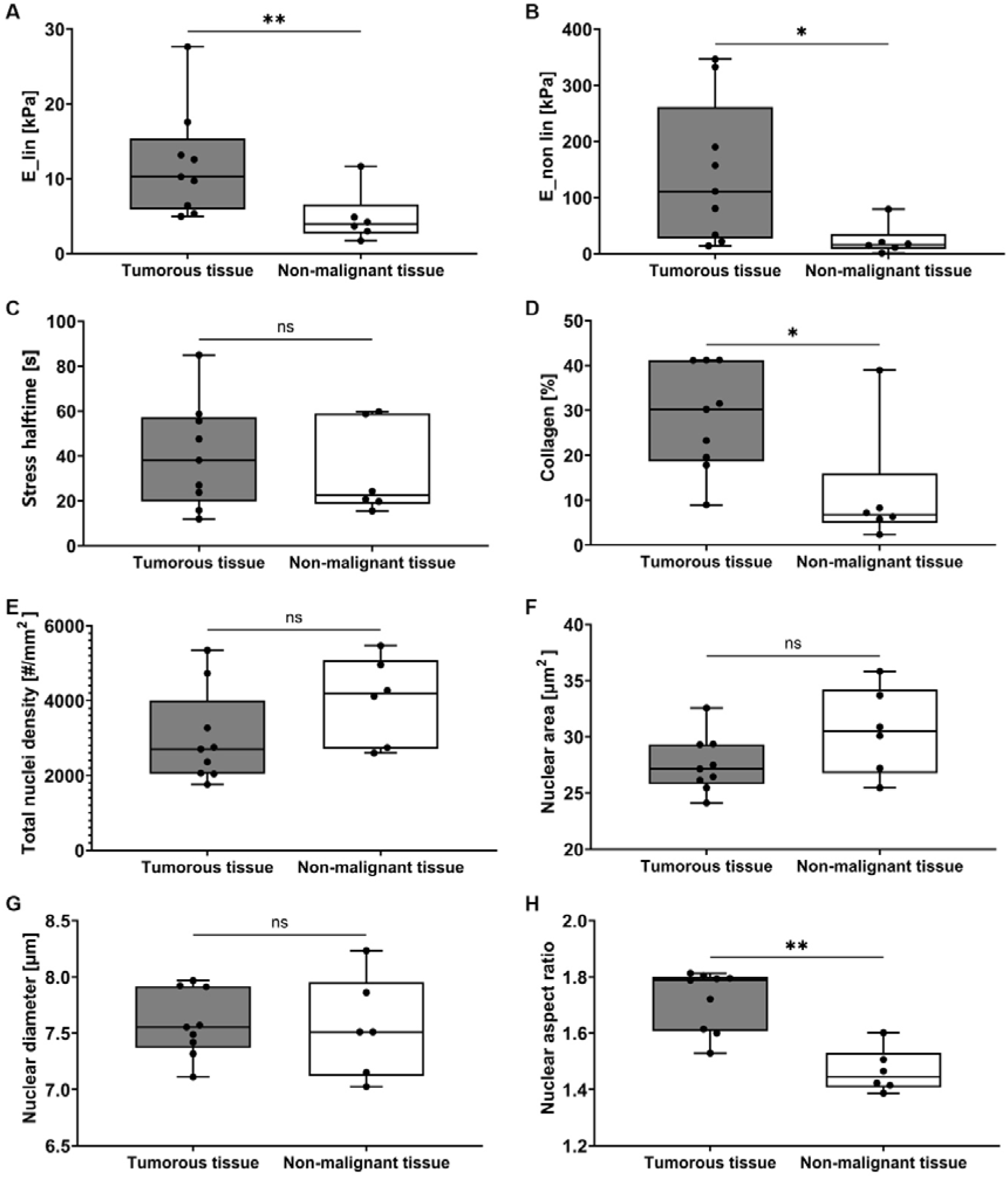
Quantification of ex vivo tissue biopsies comparing tumor tissue with adjacent non-malignant tissue: elastic modulus (A) E_lin in linear range and (B) E_non_linear in non-linear range, (C) stress relaxation halftime τ_1/2_, (D) collagen fibrous tissue area, (E) total nuclear cell density, single cell (F) nuclear area, (G) diameter and (H) aspect ratio.

From the stress vs. time curves, the stress relaxation halftime, *τ__½_*, was obtained as the time for the stress to reach half of the peak stress at constant 15% strain (Fig. 3B, Fig. 4B). In this case, no statistical difference (p = 0.689) was found between tumor samples compared to the adjacent non-malignant tissue (Fig. 5C).

After biomechanical testing, samples were fixed and prepared for histological analysis. Based on Picrosirius Red staining and polarized imaging, using an image analysis code (Suppl. Fig. S2), we quantified the % of collagen fibrous tissue area in tumor samples and adjacent non-malignant tissue (Fig. 3D-E, Fig. 4D-E). Collagen fibrous tissue area was significantly higher for the tumor samples compared to the adjacent non-malignant tissue, with p = 0.012, (Fig. 5D), confirming the fibrotic nature of PDAC tumor tissue.

Finally, based on H&E staining (Fig. 3C, Fig. 4C) and an image analysis tool, we quantified the total cell nuclei density, as well as single cell nuclear parameters such as the area, diameter and aspect ratio (Suppl. Fig. S1). We observed a trend for lower total cell nuclei density (Fig. 5E) for the tumor samples compared to the adjacent non-malignant tissue, consistent with increased stroma in tumor samples as depicted by histology (Fig. 3C, Fig. 4C, and analogous images in Suppl Info), although the differences were not statistically significant (p = 0.134). Regarding the single cell nuclear morphology, nuclei were significantly more elongated in the tumor samples (p = 0.002, Fig. 5H), consistent with the trend for smaller area (p-value=0.094, Fig. 5F) for a comparable diameter (Fig. 5G).

### 3.4 Correlation analysis of ex vivo human pancreatic tumor tissue mechanics, ECM architecture, cell nuclei density and morphology, and patient survival

Next, we performed correlation analysis of human pancreatic tumor tissue samples, comparing ex vivo tissue mechanics with ECM architecture, in particular the % of collagen fibrous tissue area, cell nuclei density and morphology, and patient survival. Fig. 6A-B show the Pearsońs R-value and p-value of all comparisons performed. We found a significant positive correlation between E_lin and E_non lin with p = 0.001 (Fig. 6C), indicating the related nature of these parameters. Interestingly, we found a significant positive correlation between collagen tissue area and E_non lin with p = 0.039 (Fig. 6E) but only a non-significant positive trend with E_lin (p = 0.092) (Fig. 6D).

**Figure 6:**
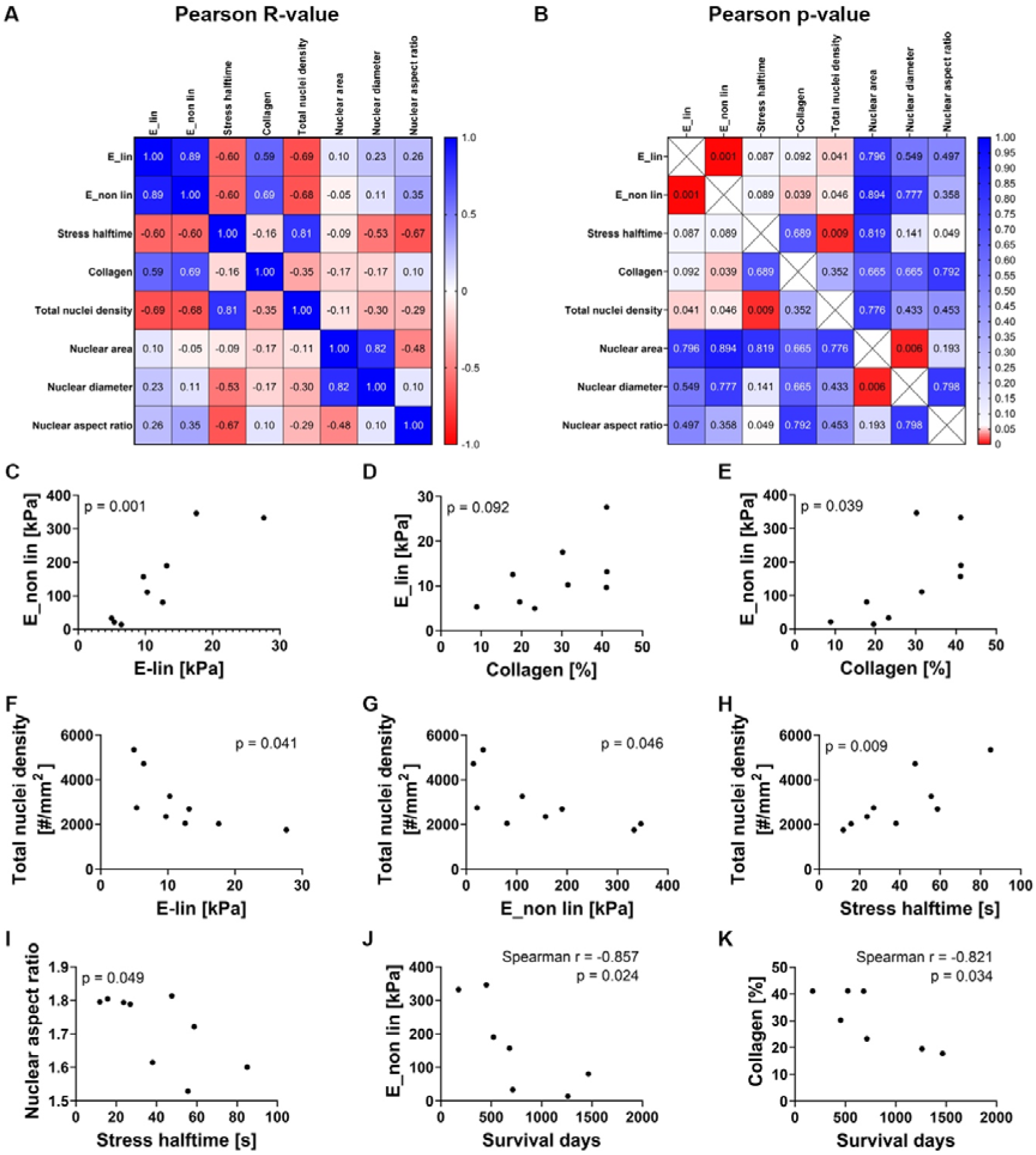
Correlation analysis of ex vivo human pancreatic tumor tissue mechanics, ECM architecture, cell nuclei density and morphology and patient survival (for patients who died). Statistical analysis: Pearsońs correlation for A-I, Spearmańs correlation for J-K.

Regarding the total cell nuclei density, we found a significant negative correlation with E_lin with p = 0.041 (Fig. 6F) and E_non lin with p = 0.046 (Fig. 6G), in contrast with a significant positive correlation with stress relaxation halftime with p = 0.009 (Fig. 6H). This suggests that the increased cellularity characteristic of healthy pancreatic tissue is correlated with a lower stiffness and higher stress relaxation halftime (less viscous). As opposed to a decreased cellularity replaced by stromal tissue in PDAC, correlated with higher stiffness and lower stress relaxation halftime (more viscous). Regarding single cell nuclear morphology, we found negative trend between single cell nuclear aspect ratio and stress relaxation halftime, associating elongated nuclei with a more viscous tumor tissue (Fig. 6I). Finally, we found a significant positive correlation between single cell nuclear area vs. diameter with p = 0.006, as expected.

Finally, we performed a Spearman correlation analysis of patient survival days for patients who died, with all ex vivo parameters. Interestingly, we found a significant negative correlation of survival time vs. collagen percentage with p = 0.024 (Fig. 6J) and vs. E_non lin with p = 0.034 (Fig. 6K). This indicates that a fibrous PDAC biopsy with increased stiffness correlates with poorer patient survival.

### 3.5 Correlation analysis of pre-operative in vivo clinical MRE data, ex vivo tissue mechanics, ECM architecture, cell nuclei density and morphology, and patient survival

Next, we performed correlation analysis of pre-operative in vivo clinical MRE data and ex vivo tissue mechanics, ECM architecture, cell nuclei density and morphology, and patient survival. Fig. 7A-B show the Pearsońs R-value and p-value of all comparisons performed. The main parameters that showed a significant correlation were the shear wave speed, c (as proxy for stiffness) and the loss angle of the complex shear modulus, (*φ* (proxy of the relative viscosity or tissue fluidity).

**Figure 7:**
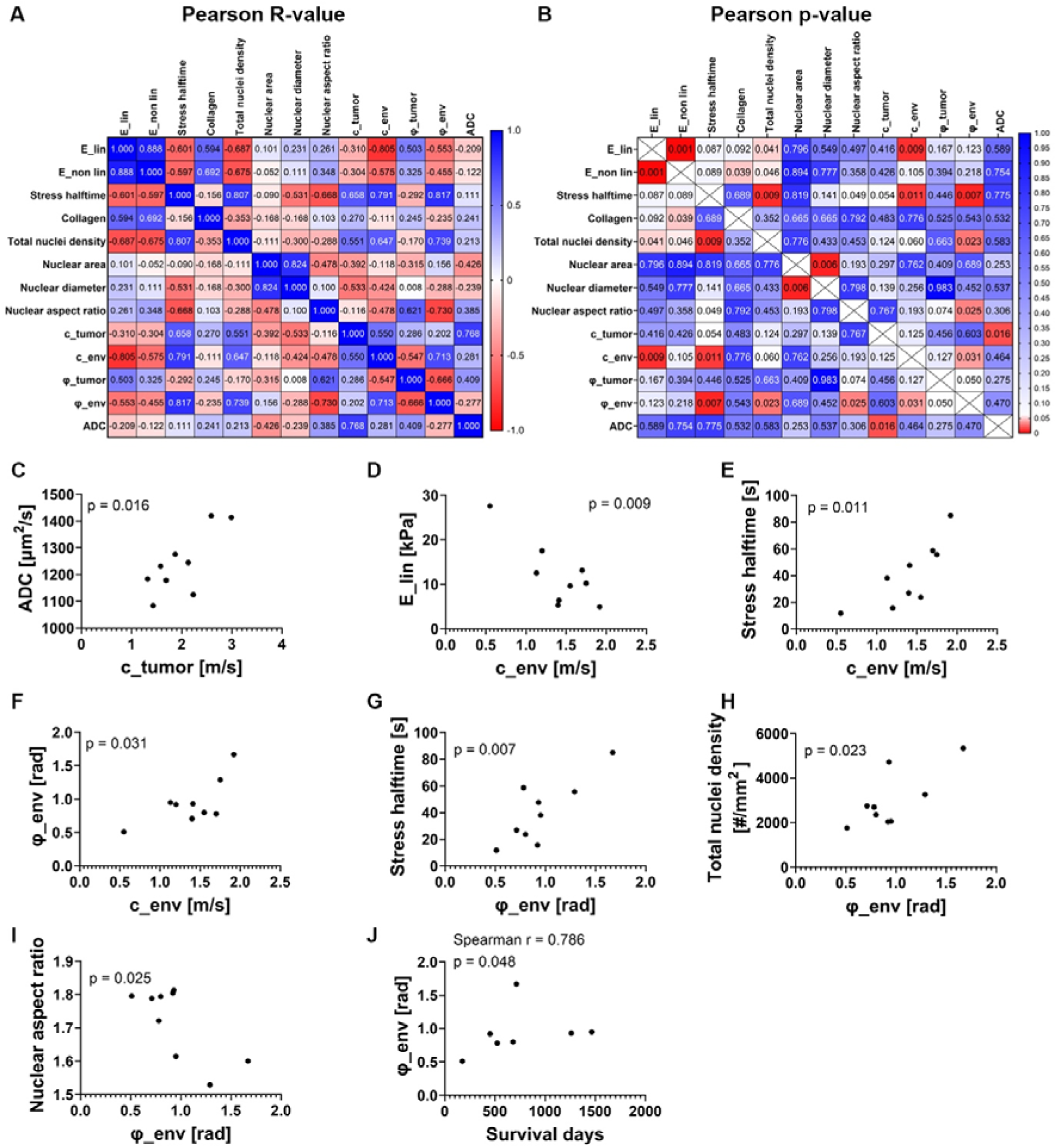
Correlation analysis of pre-operative in vivo clinical MRE data and ex vivo tissue mechanics, ECM architecture, cell nuclei density and morphology of tumor tissue. Statistical analysis: Pearsońs correlation for A-I, Spearmańs correlation for J.

Regarding the MRE properties of the tumor tissue, we found a significant positive correlation between c_tumor and ADC with p = 0.016 (Fig. 7C). With respect to the surrounding environment (env), we found a significant negative correlation between c_env and E_lin of ex vivo tumor tissue with p = 0.009 (Fig. 7D), while a significant positive correlation was found between c_env and stress relaxation halftime of ex vivo tumor tissue (lower *τ*_1/2_ indicates more viscous), with p = 0.011 (Fig. 7E).

Concerning (*φ*, we found a significant positive correlation between *ϕ*_env and c_env with p = 0.031 (Fig. 7F). Moreover, we found a significant positive correlation between *ϕ*_env and stress relaxation halftime of ex vivo tumor tissue with p = 0.007 (Fig. 7G), a significant positive correlation between *ϕ*_env and total nuclei density of ex vivo tumor tissue with p = 0.023 (Fig. 7H), and a significant negative correlation between *ϕ*_env and nuclear aspect ratio of ex vivo tumor tissue with p = 0.025 (Fig. 7I).

Following a Spearman correlation analysis of patient survival days for patients who died with MRE data, we found a significant positive correlation between *ϕ*_env and survival time with p = 0.048 (Fig. 7J). This indicates that a less viscous environment associated with poorer survival.

## 4. Discussion

In this study we evaluate extracellular matrix mechanics, tissue architecture and cellularity in human pancreatic ductal adenocarcinoma. For this we quantify, on one hand, pre-operative in vivo clinical MRE mechanical data of intact (uncut) tumor vs. surrounding environment (Fig. 2); and on the other hand, ex vivo tissue biopsy mechanical properties, tissue architecture and cellularity of tumor tissue vs. adjacent non-malignant tissue (Fig. 3-5). Finally, we perform correlation analyses of ex vivo tumor tissue mechanics, ECM architecture, cell nuclei density and morphology (Fig. 6); which is then complemented with MRE data (Fig. 7) and clinical data such as patient survival time. Fig. 8 summarizes the outcomes of the main parameters measured.

**Figure 8:**
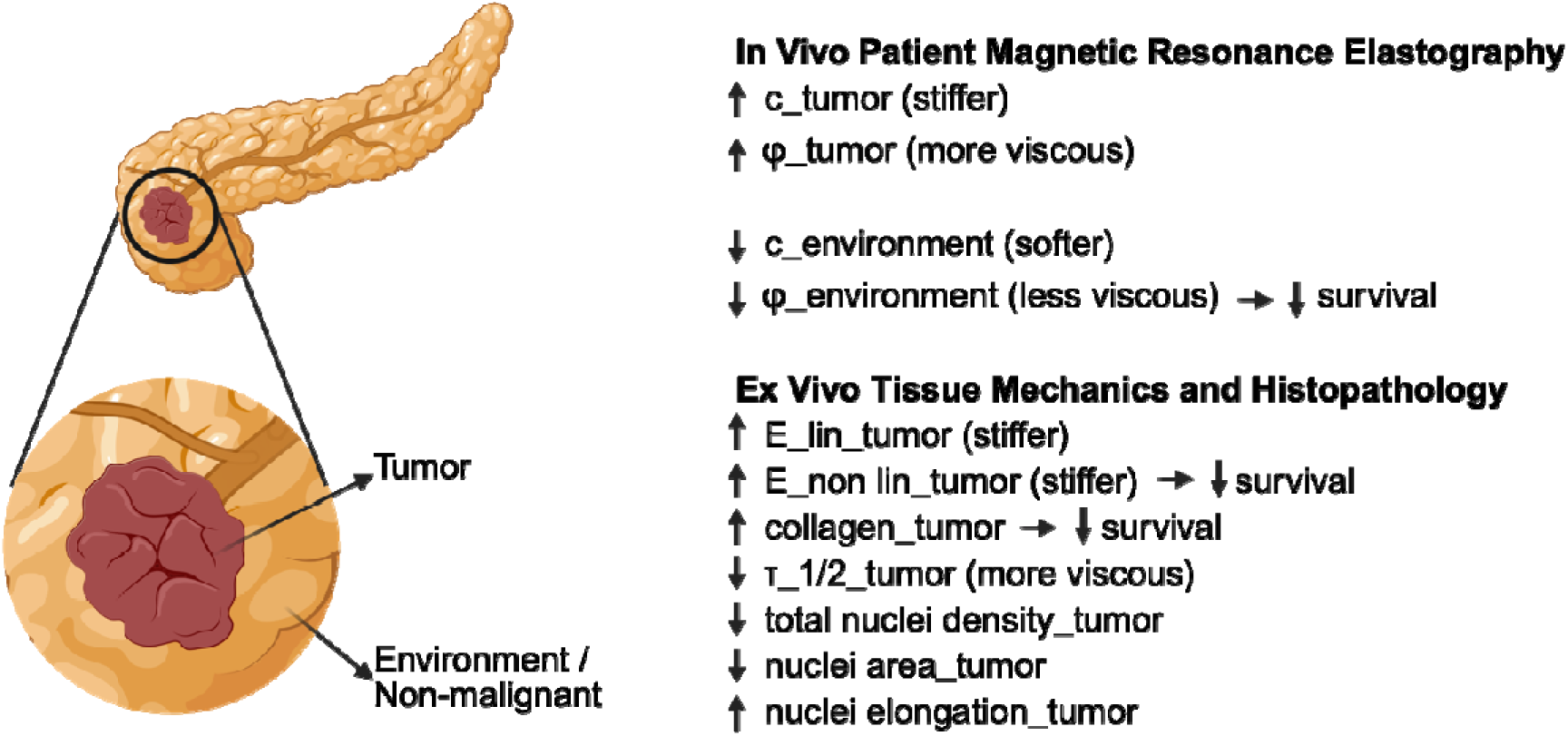
Graphical summary of the main parameters measured with MRE, ex vivo tissue mechanics and histopathology, in relation to tumor vs. environment/non-malignant. Created with Biorender.

In the analysis of the ex vivo tissue biopsies, we observe significantly higher tissue stiffness, with increased E_lin and E_non lin for tumor tissue vs. non-malignant, in agreement with previous data [4–8,10–12]. This is associated with significantly higher collagen percentage in the tumor tissue vs. non-malignant. In the correlation analysis of ex vivo tumor samples we found that E_lin and E_non lin are positively correlated, indicating the related nature of these parameters. With regard to the collagen content, there was a trend for positive correlation with E_lin and a significant positive correlation with E_non lin. This data confirms that fibrotic PDAC tumor tissue is also stiffer. Notably, none of these patients received neoadjuvant therapy, therefore this fibrotic nature is related to the disease and not to any previous treatment. Interestingly, both E_non lin and collagen content exhibit a significant negative correlation with survival days; thus, a stiffer and fibrotic tumor is indicative of poor prognosis.

In relation to total cell nuclei density, the median for tumor tissue was lower than for non-malignant tissue, confirming the replacement of a highly cellular healthy pancreatic tissue by increased stroma, although the differences were not statistically significant. In the correlation analysis of ex vivo tumor samples, this trend was confirmed with a significant negative correlation between total cell nuclei density and E_lin and E_non lin, showing that the stiffer the tumor tissue, the lower the total cell nuclei number.

Regarding the ex vivo stress relaxation halftime (lower *τ*_1/2_, more viscous), no significant differences were found between tumor tissue vs. non-malignant. However, in the correlation analysis of ex vivo tumor samples, we found a significant positive correlation between total cell nuclei density and stress relaxation halftime, suggesting that reduced cellularity associated with malignancy, corresponds to lower stress relaxation halftime (more viscous). This is in agreement with previous reported data, with stiffer and more viscous tissue for cancerous tissue [4–8,10–12].

With regard to single cell morphology, significantly more elongated nuclei and a trend for smaller nuclear area were found in the tumor tissue compared to non-malignant, which together with a lower total cell nuclei density have been associated with a state of cell unjamming and an invasive phenotype [11,29,30]. Finally, in the correlation analysis of ex vivo tumor samples a trend for a negative correlation was found for nuclear aspect ratio and stress relaxation halftime, suggesting that more elongated nuclei associated with malignancy, is also related to a lower stress relaxation halftime and more viscous nature.

Altogether, our combined image analysis of histological sections of H&E for cell density and morphology, combined with histological sections of Picrosirius red for collagen content suggests that it may be possible to extrapolate information based on H&E cell density and morphology analysis, and relate this to lower/higher stromal component and collagen fraction, associated with malignancy. This is particularly interesting, since most pathology departments can provide H&E sections, which could be analyzed by open-access image-analysis algorithms like the one used here.

Importantly, pre-operative patient in vivo clinical MRE mechanical data gives us complementary information of intact (uncut) tissue mechanics, as opposed to cut tissue explants, from both the tumor tissue and surrounding environment. Here, c is a surrogate maker of stiffness (higher c, higher stiffness); and *ϕ* is a viscosity-related parameter that measures tissue fluidity (higher *ϕ*, higher viscosity). Our reported values of higher c and *ϕ* for tumor vs. environment are in agreement with previously reported data in human pancreatic cancer [10–13]. The significant positive correlation between c_tumor with the apparent diffusion coefficient (ADC) indicates higher water diffusion in stiffer tumors. It was previously reported that higher stiffness in tumors, for instance due to collagen crosslinking, does not affect water mobility [31]. Notably, c_tumor and *ϕ*_tumor do not show any correlation with ex vivo tissue mechanics, although both methods MRE and ex vivo mechanical testing show significantly higher stiffness for tumor compared to non-malignant/environment. Furthermore, we find interesting correlations in relation to the surrounding environment (env). We observe that a softer environment (low c_env) is significantly correlated with a higher ex vivo tumor stiffness (E_lin) and lower ex vivo tumor stress relaxation halftime (more viscous), both associated with malignancy. In addition, we find a significant positive correlation between c_env and *ϕ*_env, indicating that a softer environment (low c_env) is also less viscous (low *ϕ*_env). Finally, we observed that a less viscous environment (low *ϕ*_env) is significantly correlated with lower ex vivo tumor stress relaxation halftime (more viscous), lower ex vivo tumor cell density and more elongated nuclei, all of which are associated with malignancy. This is further confirmed by the significant positive correlation between *ϕ*_env and survival time, with a less viscous environment associated with poorer survival. Interestingly, a similar trend has been reported recently in hepatocellular carcinoma (HCC), whereby liver softening began two weeks after HCC inoculation, followed by a decrease in tissue viscosity and fluidity two weeks later, while local lesions with stiffer mechanics were only detectable 5-6 weeks post injection [32]. This suggests that a softer and less viscous environment could favor the development of malignant PDAC and precede the detection of local tumor growth. Altogether, this points toward the importance of the mechanical properties of the intact (uncut) surrounding environment, accessible with quantitative MRE imaging, and how this can be related to ex vivo tumor properties and malignancy.

Due to the intricate experimental design involving patient pre-operative imaging prior to surgery, this study was performed as an exploratory hypothesis-generating pilot study. Thus, one limitation of the study is the low patient number. In addition, the work involves subjective delimitation of tumor vs. environment in MRE. Finally, ex vivo samples may not always reflect a precise distinction of tumor vs. non-malignant, as such decisions are taken in situ during the surgery, and the non-malignant resection margin might not always be entirely free of cancer-modified ECM.

## 5. Conclusions

In conclusion, ex vivo tumor tissue biopsy analysis suggests the following features as a malignant signature and lower survival: high tumor stiffness (E_lin, E_non lin), high collagen fraction, low total cell nuclei density, low nuclear area and elongated aspect ratio, and indirectly, low stress relaxation halftime (more viscous). Pre-operative in vivo clinical MRE mechanical data of intact (uncut) surrounding environment could capture a correlation between a softer (low c_env) and less viscous (low *ϕ*_env) surrounding environment with this malignant signature and lower survival. These findings support the potential of using in vivo clinical MRE as a non-invasive imaging approach to assess mechanical alterations before PDAC surgery.

## 6. Acknowledgements

This work was supported by the Einstein Center Regenerative Therapies (ECRT) Kickbox Seed Grant and following ECRT Research Grant (R.B.S., A.C.). O.M.I. was recipient of a Postdoctoral grant from the Fundación Científica Asociación Española Contra el Cáncer (POSTD234723MITX) and would like to acknowledge the funding received by Gipuzkoako Foru Aldundia-Diputación Foral de Gipuzkoa (grant 2025-CIE4-000015-01). A.C. and D.S.G acknowledge the funding by the DFG Emmy Noether grant CI 203/2-1. A.C. would like to acknowledge funding from IKERBASQUE Basque Foundation for Science, from Fundación Científica Asociación Española Contra el Cáncer (grant LABAE223466CIPI), from the Spanish Ministry of Science and Innovation (MCIN/AEI/10.13039/501100011033/FEDER UE, through grant PID2021-123013OB-I00) and from the European Research Council Consolidator Grant (DORMATRIX, 101123883). We would also like to thank Anja Schirmeier for her assistance.

## 7. Author contribution

**Oihane Mitxelena-Iribarren:** Investigation, Formal analysis, Software, Validation, Visualization, Writing - Original Draft. **Daniela S. Garske:** Investigation, Formal analysis, Software. **Dag Wulsten:** Investigation, Formal analysis, Software. **Iñigo Mendizabal-Arrieta:** Software. **Kristin Spirgath:** Software. **Salma Almutawakel**: Investigation. **Rosa B. Schmuck:** Conceptualization, Resources. **Ingolf Sack:** Conceptualization, Methodology, Validation, Resources. **Amaia Cipitria:** Conceptualization, Methodology, Validation, Resources, Visualization, Writing - Original Draft, Supervision, Project administration, Funding acquisition. All authors reviewed and edited the last version of the manuscript.

## 8. Data availability

All data supporting this article and analysis code are available at Zenodo repository (10.5281/zenodo.18172042).

## References

[1] Rahib L, Smith BD, Aizenberg R, Rosenzweig AB, Fleshman JM, Matrisian LM. Projecting Cancer Incidence and Deaths to 2030: The Unexpected Burden of Thyroid, Liver, and Pancreas Cancers in the United States. Cancer Res 2014;74:2913–21. 10.1158/0008-5472.CAN-14-0155.

[2] Bengtsson A, Andersson R, Ansari D. The actual 5-year survivors of pancreatic ductal adenocarcinoma based on real-world data. Sci Rep 2020;10:16425. 10.1038/s41598-020-73525-y.

[3] Sherman MH, Beatty GL. Tumor Microenvironment in Pancreatic Cancer Pathogenesis and Therapeutic Resistance. Annual Review of Pathology: Mechanisms of Disease 2023;18:123–48. 10.1146/annurev-pathmechdis-031621-024600.

[4] Kpeglo D, Hughes MDG, Dougan L, Haddrick M, Knowles MA, Evans SD, et al. Modeling the mechanical stiffness of pancreatic ductal adenocarcinoma. Matrix Biol Plus 2022;14:100109. 10.1016/j.mbplus.2022.100109.

[5] Rice AJ, Cortes E, Lachowski D, Cheung BCH, Karim SA, Morton JP, et al. Matrix stiffness induces epithelial–mesenchymal transition and promotes chemoresistance in pancreatic cancer cells. Oncogenesis 2017;6:e352–e352. 10.1038/oncsis.2017.54.

[6] Rubiano A, Delitto D, Han S, Gerber M, Galitz C, Trevino J, et al. Viscoelastic properties of human pancreatic tumors and in vitro constructs to mimic mechanical properties. Acta Biomater 2018;67:331–40. 10.1016/j.actbio.2017.11.037.

[7] Massey A, Stewart J, Smith C, Parvini C, McCormick M, Do K, et al. Mechanical properties of human tumour tissues and their implications for cancer development. Nature Reviews Physics 2024;6:269–82. 10.1038/s42254-024-00707-2.

[8] Peerani E, Candido JB, Maniati E, Tomás-Bort E, Sharma S, Clegg J, et al. Matrix stiffness shapes transcriptional profiles and drug responses of pancreatic cancer cells. Acta Biomater 2026;210:67–81. 10.1016/j.actbio.2025.12.020.

[9] Sack I. Magnetic resonance elastography from fundamental soft-tissue mechanics to diagnostic imaging. Nature Reviews Physics 2022;5:25–42. 10.1038/s42254-022-00543-2.

[10] Zhu L, Guo J, Jin Z, Xue H, Dai M, Zhang W, et al. Distinguishing pancreatic cancer and autoimmune pancreatitis with in vivo tomoelastography. Eur Radiol 2021;31:3366–74. 10.1007/s00330-020-07420-5.

[11] Sauer F, Grosser S, Shahryari M, Hayn A, Guo J, Braun J, et al. Changes in Tissue Fluidity Predict Tumor Aggressiveness In Vivo. Advanced Science 2023;10:2303523. 10.1002/advs.202303523.

[12] Marticorena Garcia SR, Zhu L, Gültekin E, Schmuck R, Burkhardt C, Bahra M, et al. Tomoelastography for Measurement of Tumor Volume Related to Tissue Stiffness in Pancreatic Ductal Adenocarcinomas. Invest Radiol 2020;55:769–74. 10.1097/RLI.0000000000000704.

[13] Schattenfroh J, Almutawakel S, Bieling J, Castelein J, Estrella M, Garteiser P, et al. Technical Recommendation on Multi-Driver Multifrequency MR Elastography for Tomographic Mapping of Abdominal Stiffness With a Focus on the Pancreas and Pancreatic Ductal Adenocarcinoma. Invest Radiol 2026;61:326–33. 10.1097/RLI.0000000000001231.

[14] Klein F, Sauer IM, Pratschke J, Bahra M. Bovine Serum Albumin-Glutaraldehyde Sealed Fish-Mouth Closure of the Pancreatic Remnant during Distal Pancreatectomy. HPB Surgery 2017;2017:1–7. 10.1155/2017/9747421.

[15] Schmuck RB, Brokat C, Andreou A, Felsenstein M, Klein F, Sinn B, et al. Clinicopathological Stratification and Long-term Follow-up of Patients with Periampullary Carcinomas. Anticancer Res 2018;38:5379–86. 10.21873/anticanres.12867.

[16] Bassi C, Dervenis C, Butturini G, Fingerhut A, Yeo C, Izbicki J, et al. Postoperative pancreatic fistula: An international study group (ISGPF) definition. Surgery 2005;138:8–13. 10.1016/j.surg.2005.05.001.

[17] Schmuck RB, Lippens E, Wulsten D, Garske DS, Strönisch A, Pratschke J, et al. Role of extracellular matrix structural components and tissue mechanics in the development of postoperative pancreatic fistula. J Biomech 2021;128:110714. 10.1016/j.jbiomech.2021.110714.

[18] Dittmann F, Hirsch S, Tzschätzsch H, Guo J, Braun J, Sack I. In vivo wideband multifrequency MR elastography of the human brain and liver. Magn Reson Med 2016;76:1116–26. 10.1002/mrm.26006.

[19] Gandhi D, Kalra P, Raterman B, Mo X, Dong H, Kolipaka A. Magnetic Resonance Elastography of kidneys: SELJEPI MRE reproducibility and its comparison to GRE MRE. NMR Biomed 2019;32:e4141. 10.1002/nbm.4141.

[20] Meyer T, Marticorena Garcia S, Tzschätzsch H, Herthum H, Shahryari M, Stencel L, et al. Comparison of inversion methods in <scp>MR</scp> elastography: An openLJaccess pipeline for processing multifrequency shear wave data and demonstration in a phantom, human kidneys, and brain. Magn Reson Med 2022;88:1840–50. 10.1002/mrm.29320.

[21] Tzschätzsch H, Guo J, Dittmann F, Hirsch S, Barnhill E, Jöhrens K, et al. Tomoelastography by multifrequency wave number recovery from time-harmonic propagating shear waves. Med Image Anal 2016;30:1–10. 10.1016/j.media.2016.01.001.

[22] Streitberger K-J, Reiss-Zimmermann M, Freimann FB, Bayerl S, Guo J, Arlt F, et al. High-Resolution Mechanical Imaging of Glioblastoma by Multifrequency Magnetic Resonance Elastography. PLoS One 2014;9:e110588. 10.1371/journal.pone.0110588.

[23] Streitberger K-J, Lilaj L, Schrank F, Braun J, Hoffmann K-T, Reiss-Zimmermann M, et al. How tissue fluidity influences brain tumor progression. Proceedings of the National Academy of Sciences 2020;117:128–34. 10.1073/pnas.1913511116.

[24] Schmidt U, Weigert M, Broaddus C, Myers G. Cell Detection with Star-Convex Polygons. Lecture Notes in Computer Science (including subseries Lecture Notes in Artificial Intelligence and Lecture Notes in Bioinformatics), vol. 11071 LNCS, Springer Verlag; 2018, p. 265–73. 10.1007/978-3-030-00934-2_30.

[25] Weigert M, Schmidt U. Nuclei Instance Segmentation and Classification in Histopathology Images with Stardist. 2022 IEEE International Symposium on Biomedical Imaging Challenges (ISBIC), IEEE; 2022, p. 1–4. 10.1109/ISBIC56247.2022.9854534.

[26] Weigert M, Schmidt U, Haase R, Sugawara K, Myers G. Star-convex Polyhedra for 3D Object Detection and Segmentation in Microscopy. 2020 IEEE Winter Conference on Applications of Computer Vision (WACV), IEEE; 2020, p. 3655–62. 10.1109/WACV45572.2020.9093435.

[27] Junqueira LCU, Bignolas G, Brentani RR. Picrosirius staining plus polarization microscopy, a specific method for collagen detection in tissue sections. Histochem J 1979;11:447–455. 10.1007/BF01002772.

[28] Cipitria A, Wagermaier W, Zaslansky P, Schell H, Reichert JC, Fratzl P, et al. BMP delivery complements the guiding effect of scaffold architecture without altering bone microstructure in critical-sized long bone defects: A multiscale analysis. Acta Biomater 2015;23:282–94. 10.1016/J.ACTBIO.2015.05.015.

[29] Fuhs T, Wetzel F, Fritsch AW, Li X, Stange R, Pawlizak S, et al. Rigid tumours contain soft cancer cells. Nat Phys 2022;18:1510–9. 10.1038/s41567-022-01755-0.

[30] Gottheil P, Lippoldt J, Grosser S, Renner F, Saibah M, Tschodu D, et al. State of Cell Unjamming Correlates with Distant Metastasis in Cancer Patients. Phys Rev X 2023;13:031003. 10.1103/PhysRevX.13.031003.

[31] Sauer F, Oswald L, Ariza de Schellenberger A, Tzschätzsch H, Schrank F, Fischer T, et al. Collagen networks determine viscoelastic properties of connective tissues yet do not hinder diffusion of the aqueous solvent. Soft Matter 2019;15:3055–64. 10.1039/C8SM02264J.

[32] de Moraes PAD, Safraou Y, Krehl K, Häckel A, Haase T, Metzkow S, et al. Longitudinal in vivo MR elastography reveals whole-liver viscoelastic involvement in a murine model of hepatocellular carcinoma. BioRxiv 2025:2025.09.04.674138. 10.1101/2025.09.04.674138.

