## Supplementary Information for "Extracellular Matrix Mechanobiology in Pancreatic Ductal Adenocarcinoma: Correlating In Vivo Patient Magnetic Resonance Elastography with Ex Vivo Tissue Mechanics and Histopathology"

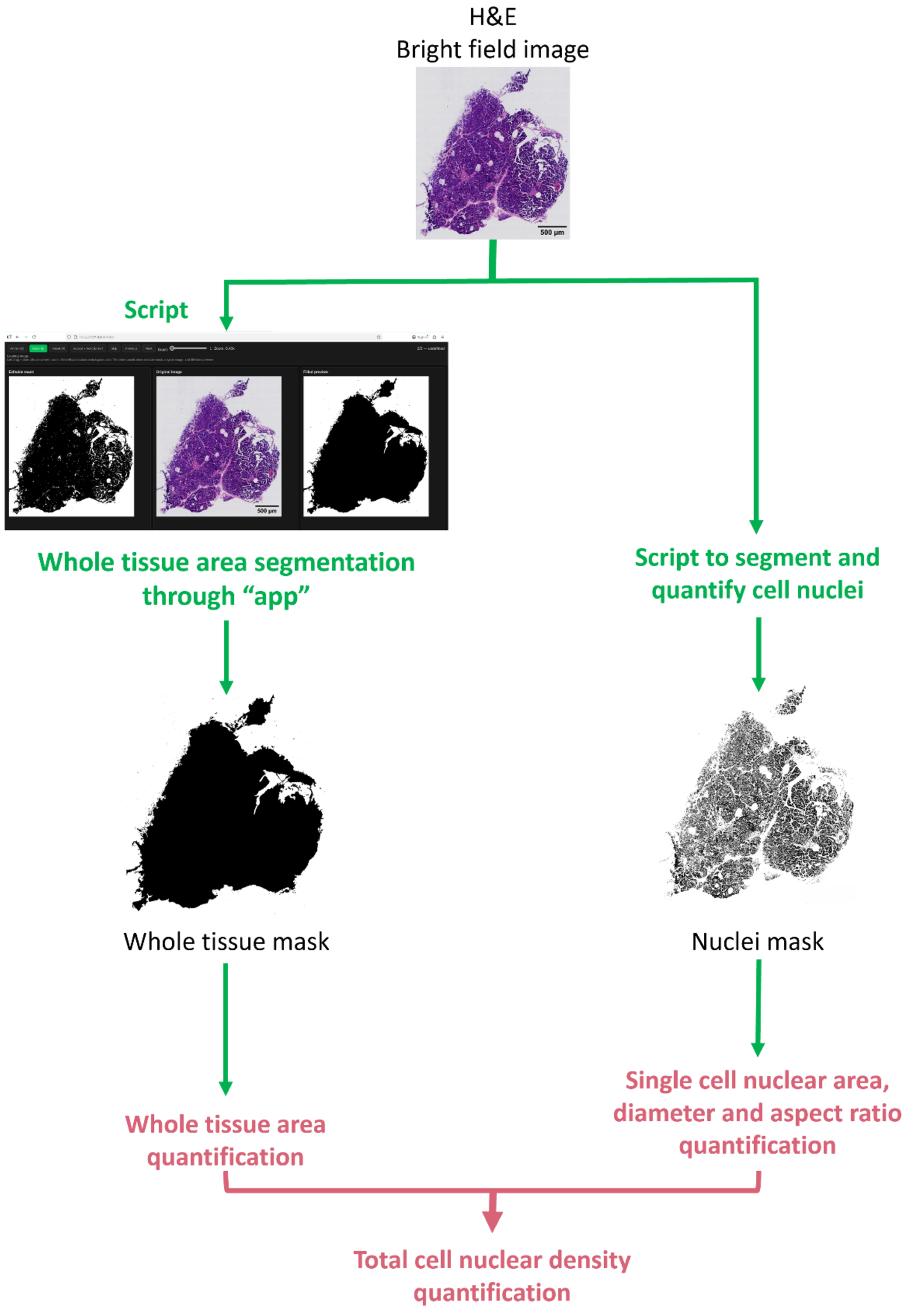


Supplementary Fig. S1: Analysis of H&E stained images to quantify cell density and cell morphological characteristics. The Python code is available in the Zenodo data repository.

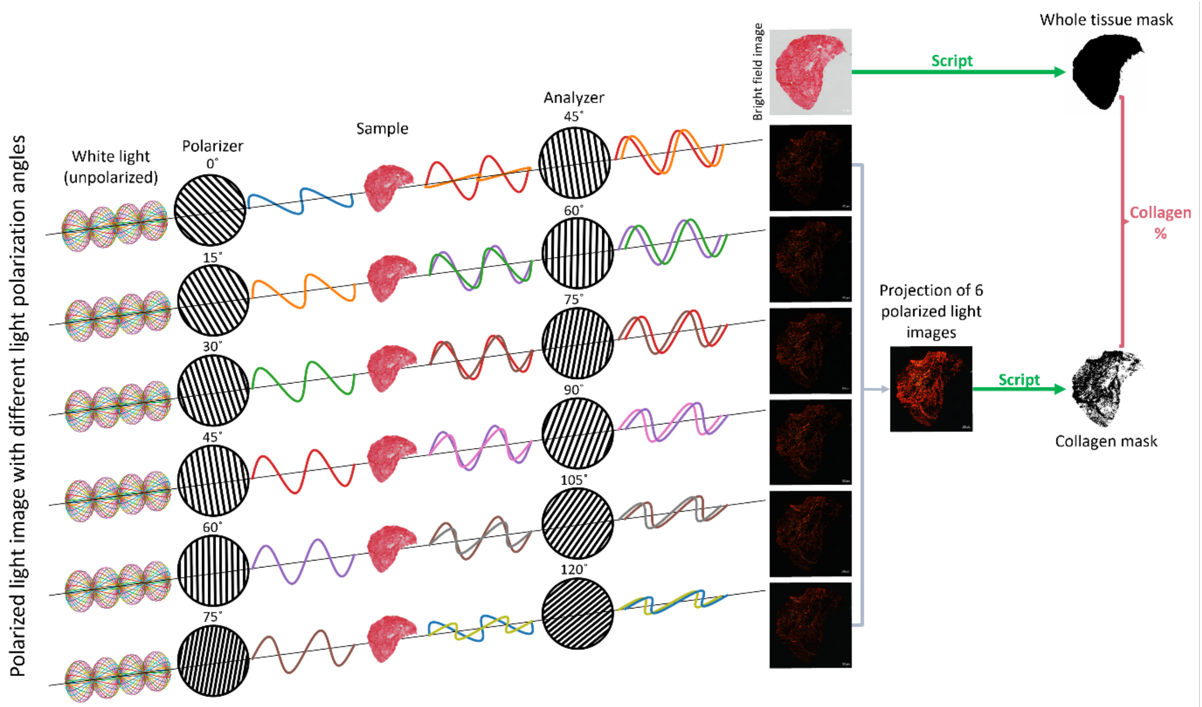


Supplementary Fig. S2: Acquisition of Picrosirius Red stained images with bright field and polarized light at different angles, and analysis to quantify the fraction of collagen fibrous tissue. For more information, see Python code in the Zenodo data repository.

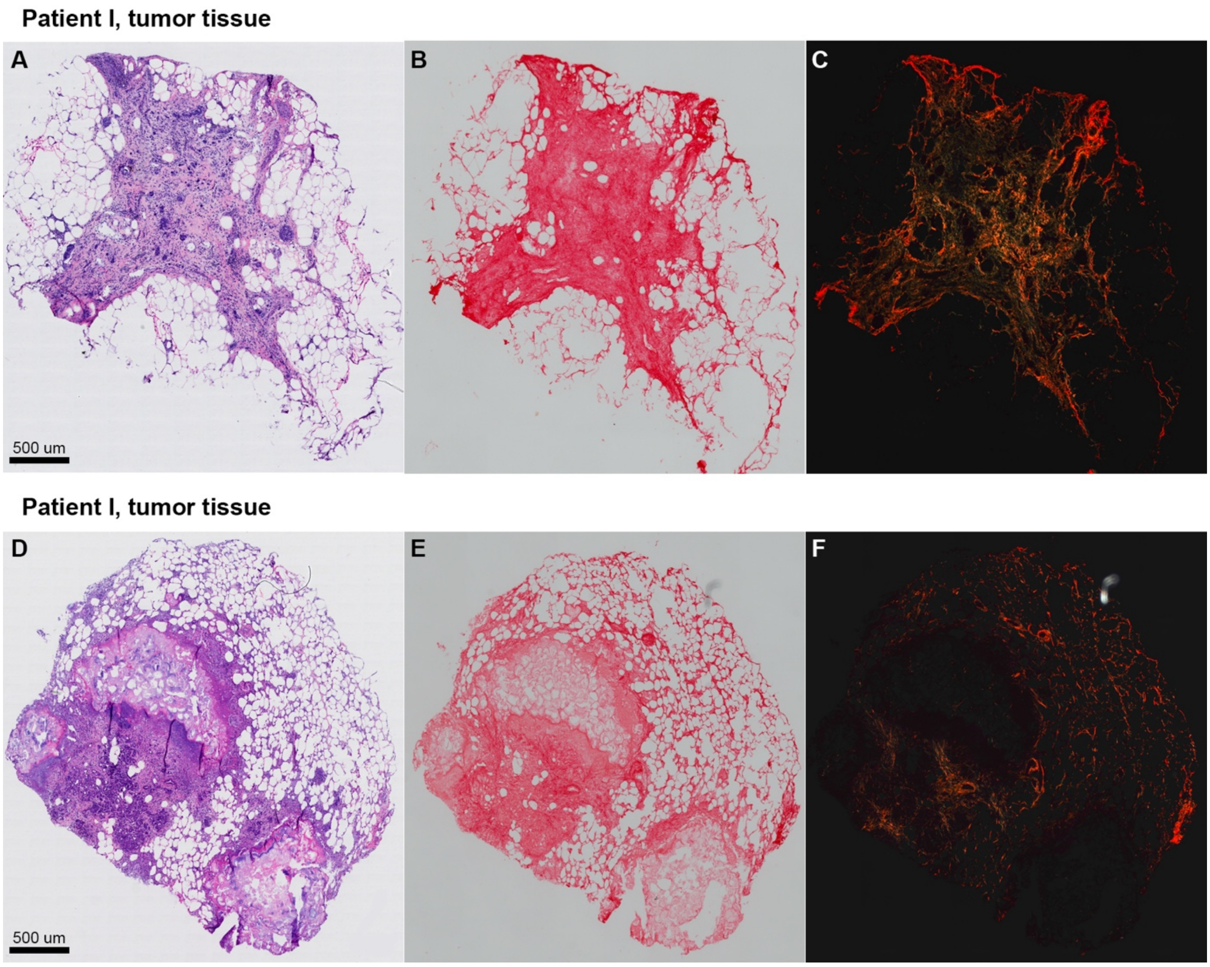


Supplementary Fig. S3: Histology of patient I, with H&E staining (A, D), Picrosirius red staining with bright field (B, E) and polarized light imaging (C, F) of tumor tissue of two different samples.

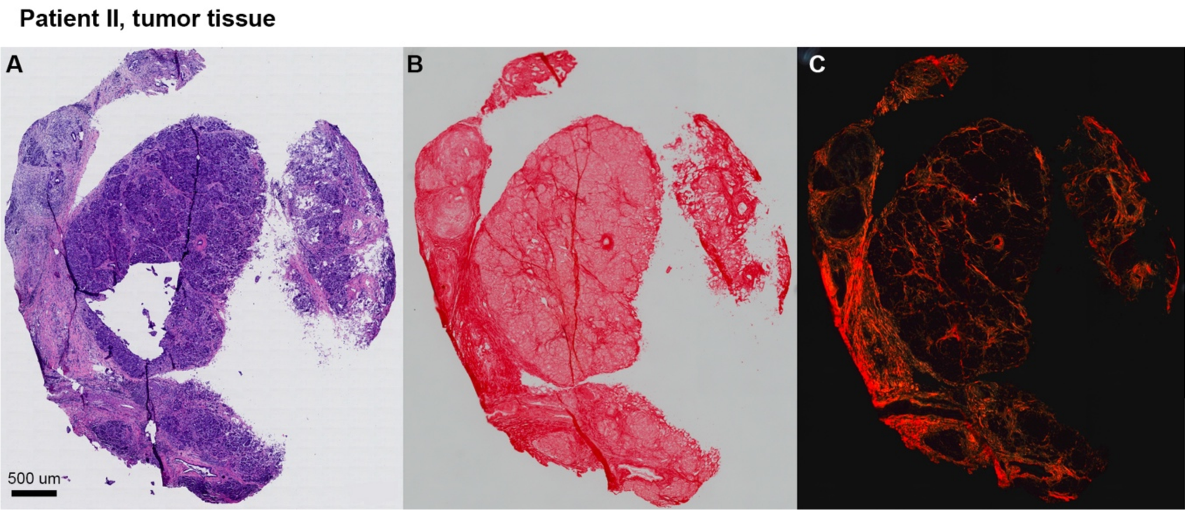


Supplementary Fig. S4: Histology of patient II, with H&E staining (A), Picrosirius red staining with bright field (B) and polarized light imaging (C) of tumor tissue.

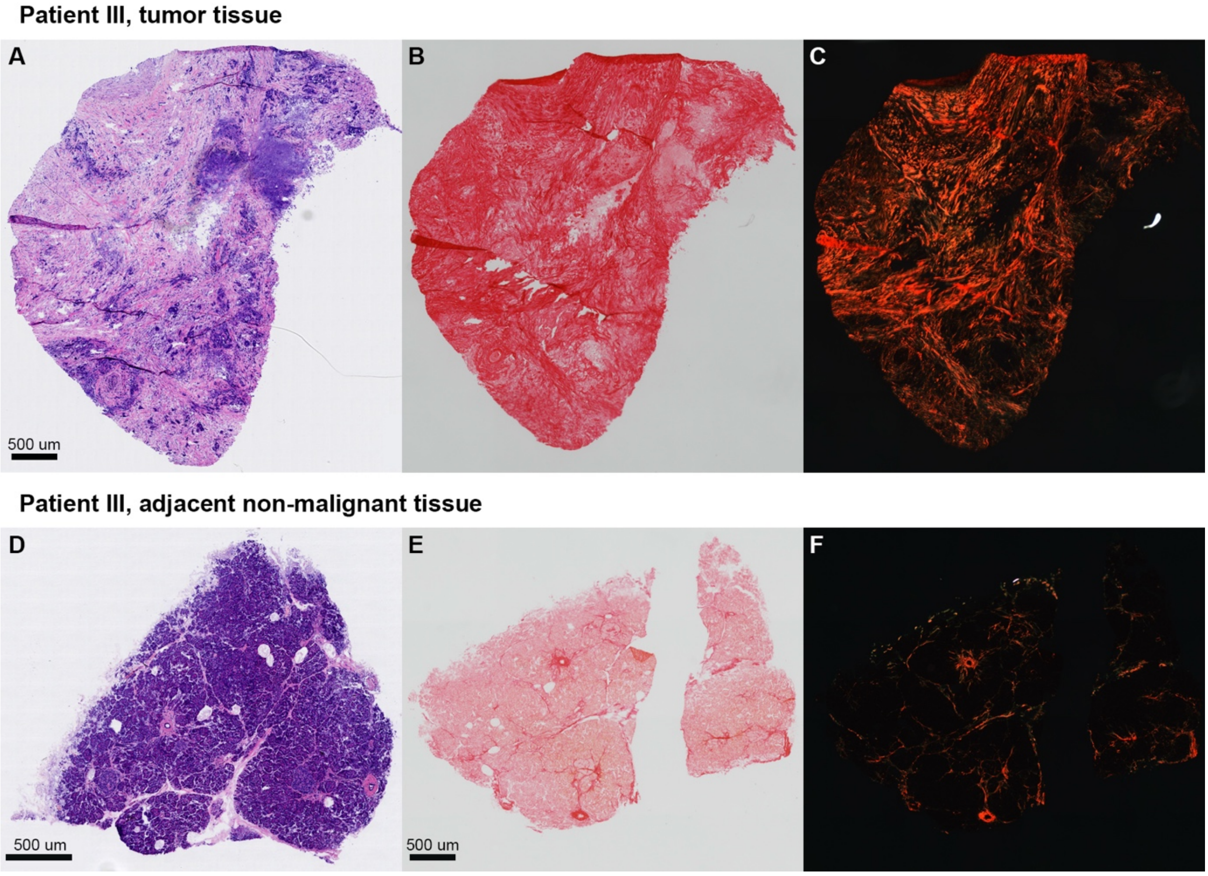


Supplementary Fig. S5: Histology of patient III, with H&E staining (A, D), Picrosirius red staining with bright field (B, E) and polarized light imaging (C, F), of tumor tissue (A-C) compared to adjacent non-malignant tissue (D-F).

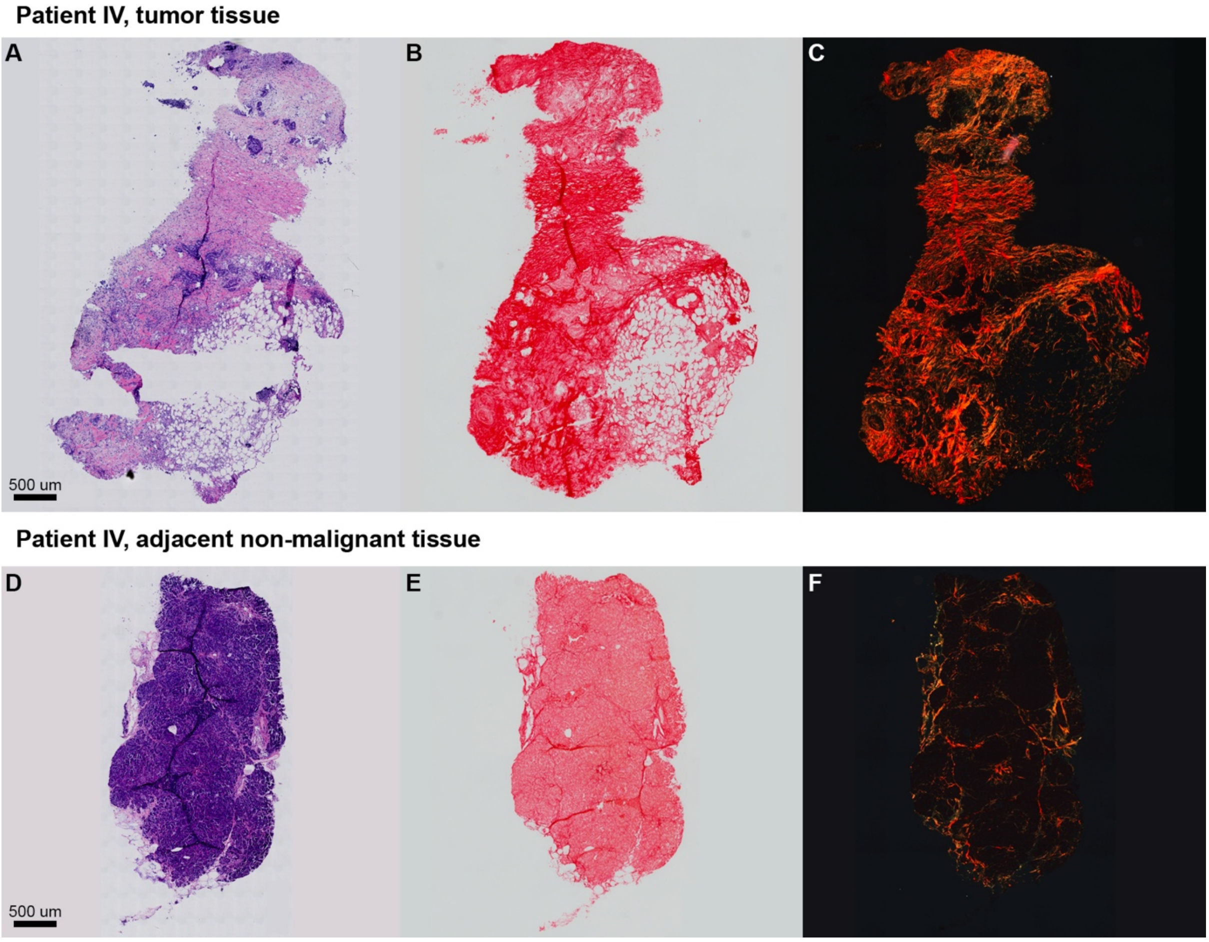


Supplementary Fig. S6: Histology of patient IV, with H&E staining (A, D), Picrosirius red staining with bright field (B, E) and polarized light imaging (C, F), of tumor tissue (A-C) compared to adjacent non-malignant tissue (D-F).

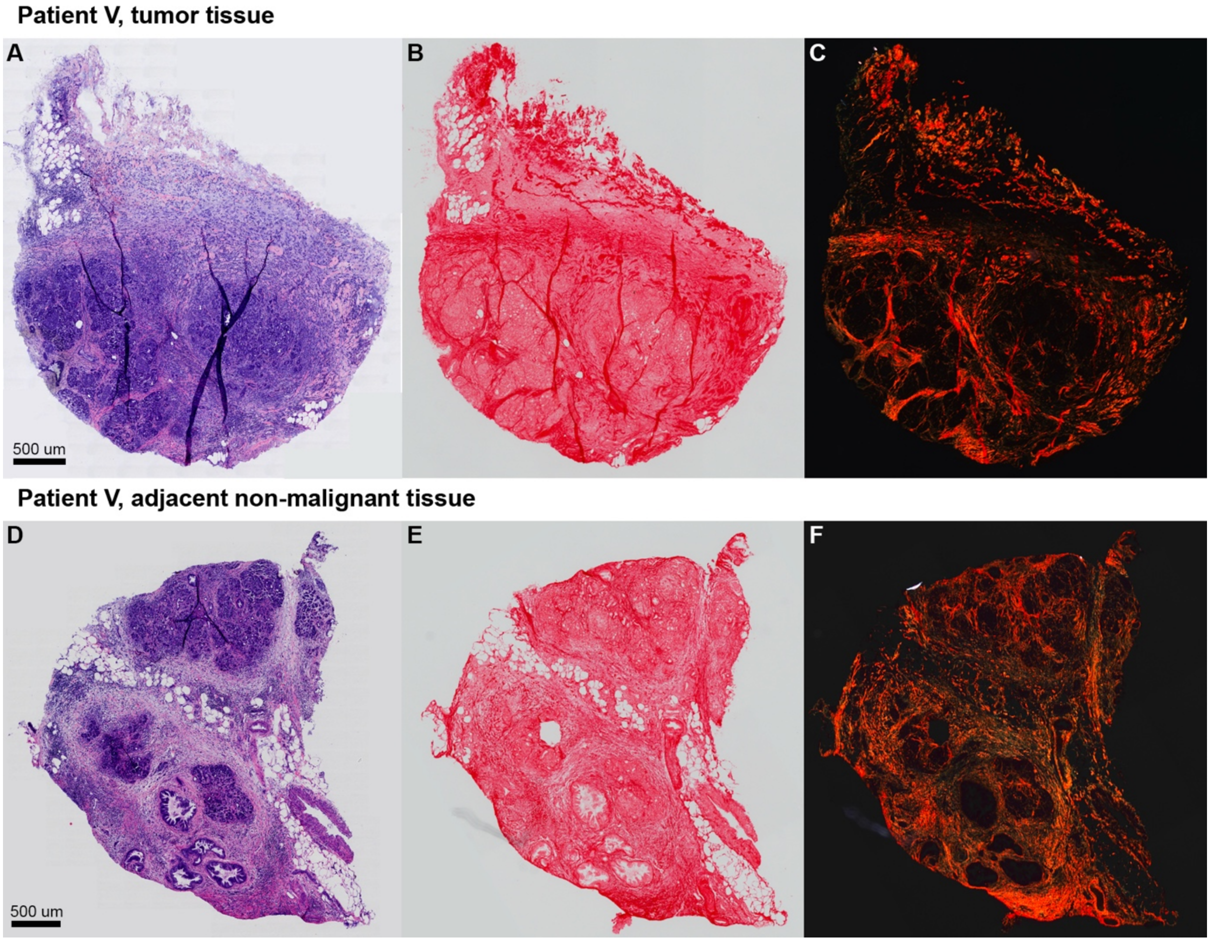

Supplementary Fig. S7: Histology of patient V, with H&E staining (A, D), Picrosirius red staining with bright field (B, E) and polarized light imaging (C, F), of tumor tissue (A-C) compared to adjacent non-malignant tissue (D-F).

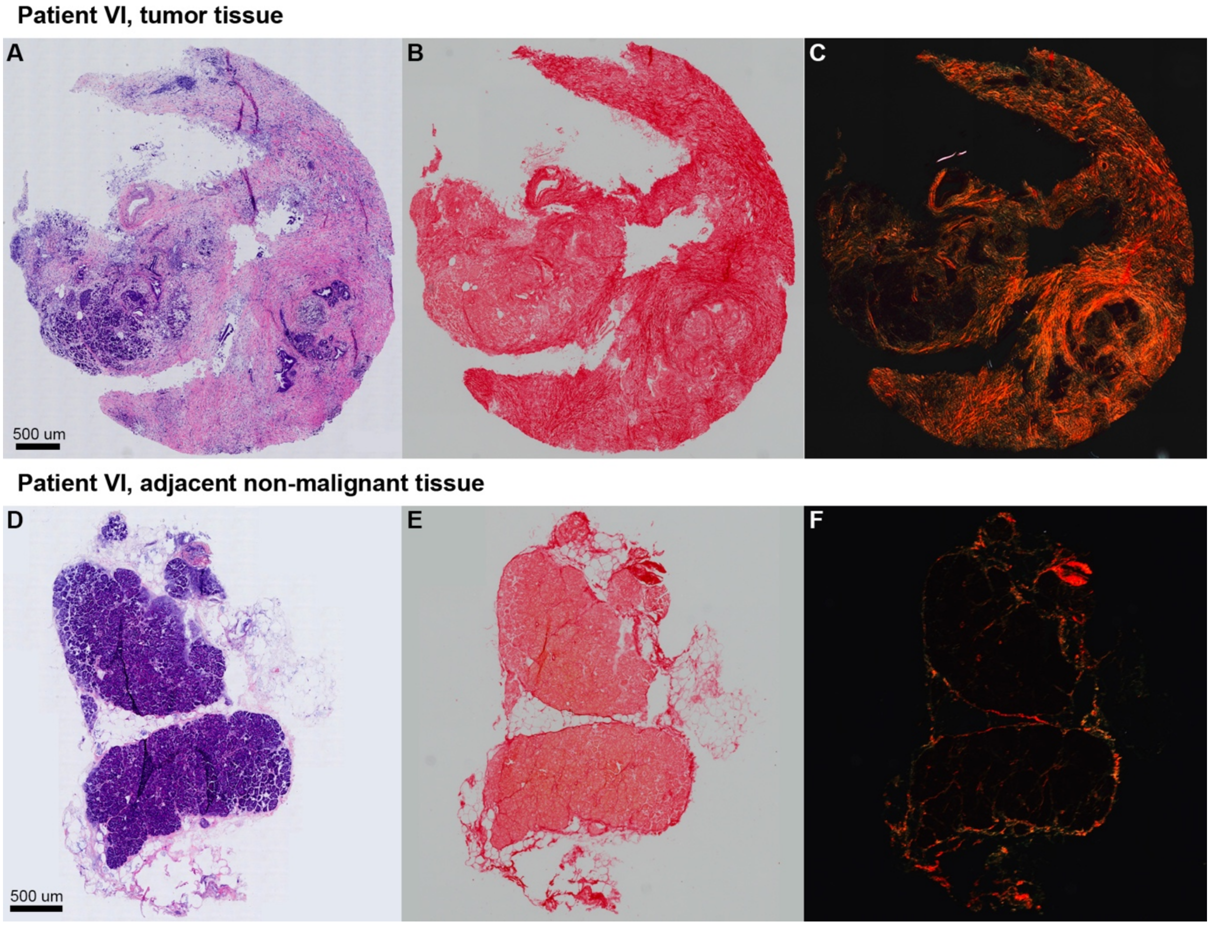

Supplementary Fig. S8: Histology of patient VI, with H&E staining (A, D), Picrosirius red staining with bright field (B, E) and polarized light imaging (C, F), of tumor tissue (A-C) compared to adjacent non-malignant tissue (D-F).

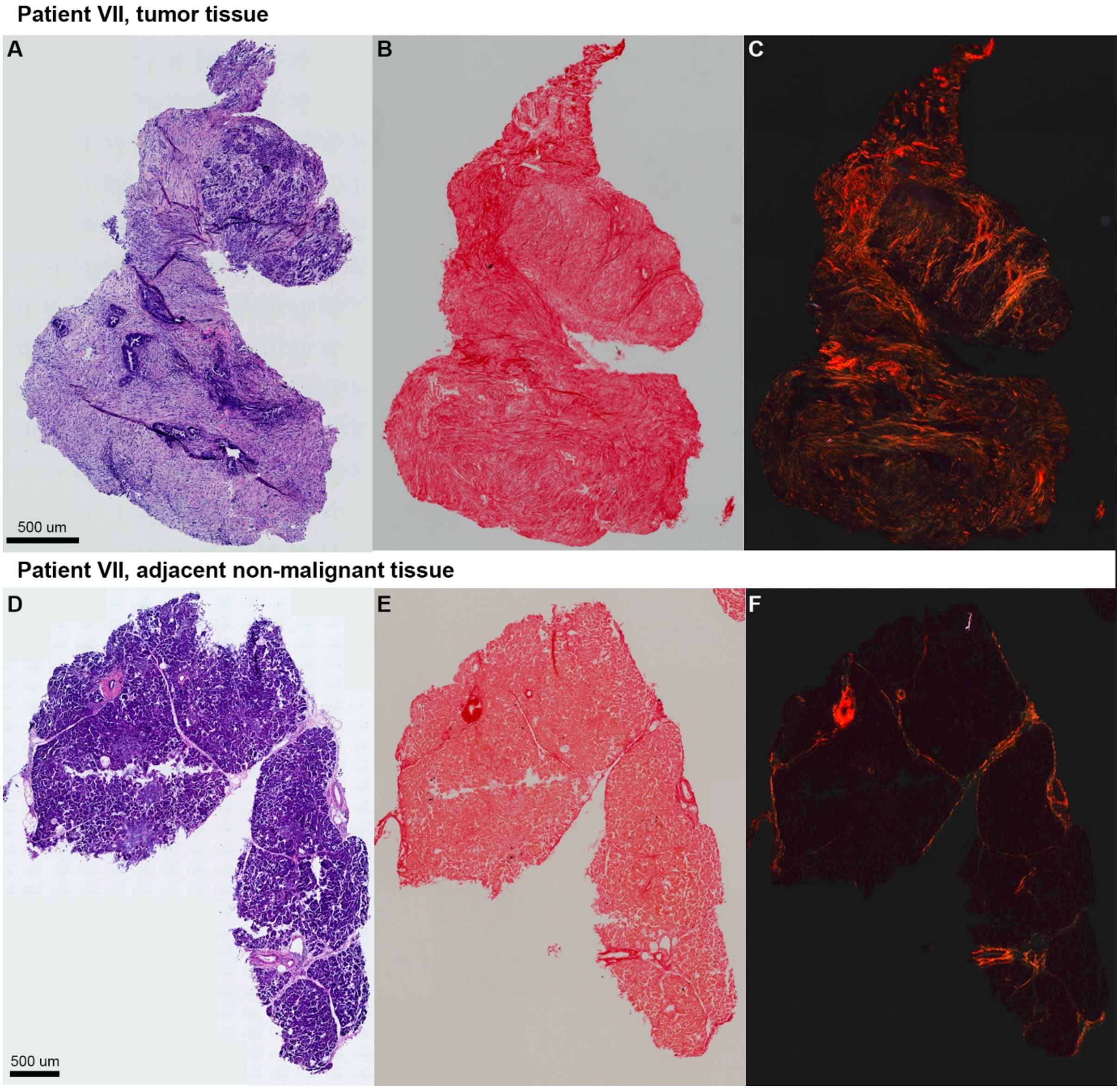

Supplementary Fig. S9: Histology of patient VII, with H&E staining (A, D), Picrosirius red staining with bright field (B, E) and polarized light imaging (C, F), of tumor tissue (A-C) compared to adjacent non-malignant tissue (D-F).

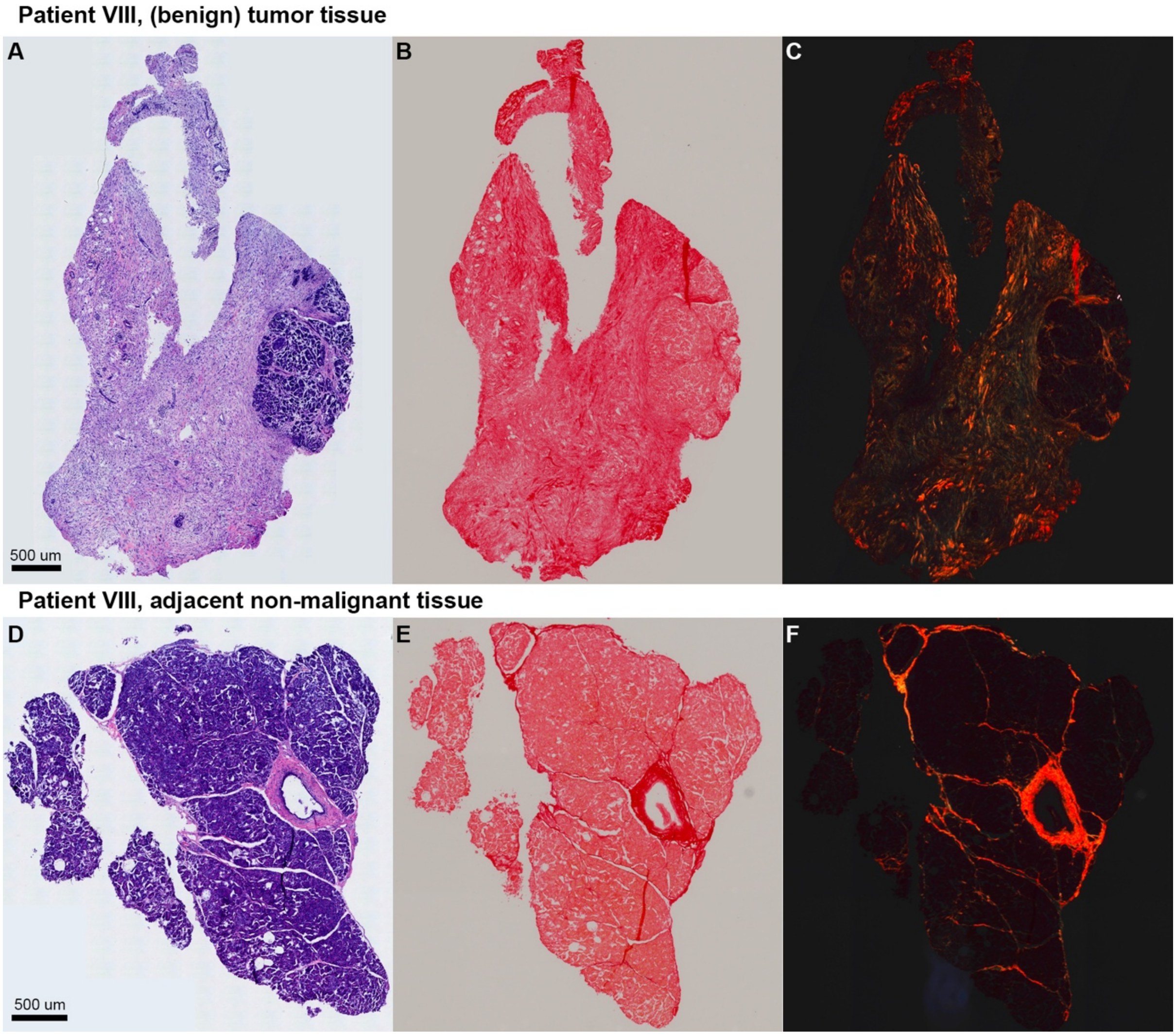


Supplementary Fig. S10: Histology of patient VIII, with H&E staining (A, D), Picrosirius red staining with bright field (B, E) and polarized light imaging (C, F), of (benign) tumor tissue (A-C) compared to adjacent non-malignant tissue (D-F).

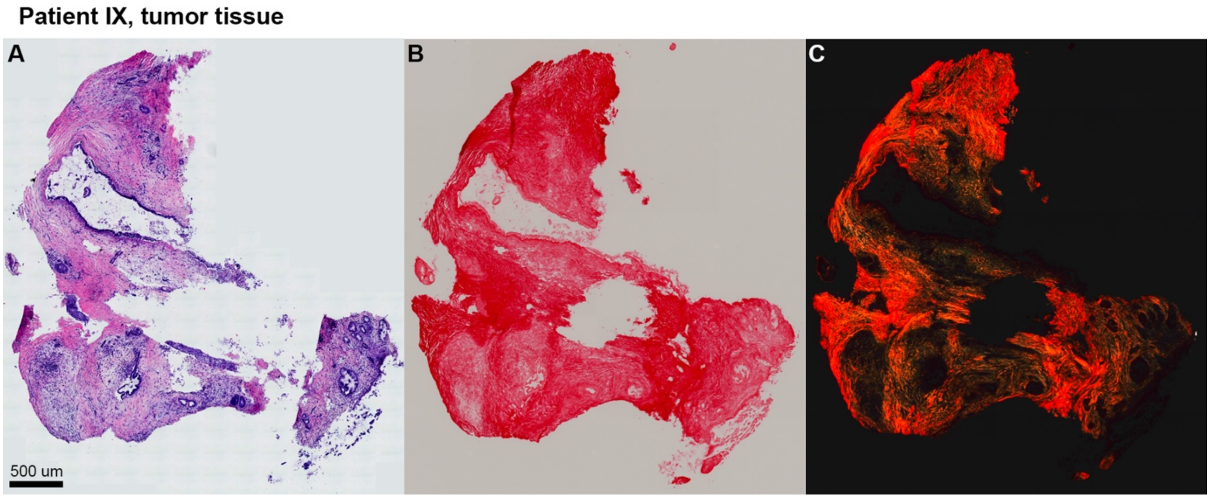


Supplementary Fig. S11: Histology of patient IX, with H&E staining (A), Picrosirius red staining with bright field (B) and polarized light imaging (C) of tumor tissue.
